# MESSI: Multimodal Experiments with SyStematic Interrogation using nextflow

**DOI:** 10.64898/2026.03.09.710452

**Authors:** Chunqing (Tony) Liang, Tajveer Grewal, Asees Singh, Amrit Singh

**Author notes:** <u>Corresponding author</u> Amrit Singh, PhD, Department of Anesthesiology, Pharmacology and Therapeutics, UBC Centre for Heart Lung Innovation, Room 166 - 1081 Burrard Street Vancouver, BC, Canada.

## Abstract

**Background:** Multimodal biomedical studies increasingly profile multiple molecular and clinical modalities from the same samples, creating new opportunities for disease prediction and biological discovery. However, benchmarking multimodal integration methods remains difficult because studies often use inconsistent preprocessing, unequal tuning strategies, and non-comparable evaluation schemes, limiting fair assessment across methods.

**Results:** We developed MESSI (Multimodal Experiments with SyStematic Interrogation), a reproducible Nextflow-based benchmarking framework for multimodal outcome prediction that standardizes data preparation, supports interoperable R and Python workflows, and enforces leakage-free nested cross-validation for model selection and model assessment. MESSI currently implements representative intermediate- and late-integration methods and supports bulk multiomics, bulk multimodal, and single-cell multiomics datasets. In simulation studies with known ground truth, most methods were well calibrated in the absence of signal and achieved high performance under strong signal, whereas differences emerged under weaker signal and in feature recovery. We then applied MESSI to 19 real datasets spanning cancer, neurodevelopmental, neurodegenerative, infectious, renal, transplant, and metastatic disease settings, with diverse modality combinations including transcriptomic, epigenomic, proteomic, imaging, electrical, clinical, and single-cell-derived features. Across bulk multimodal datasets, classification differences were generally modest, although DIABLO and multiview cooperative learning tended to rank highest, while MOFA+glmnet and MOGONET were weaker overall. Biological enrichment analyses revealed clearer differences: DIABLO, RGCCA, MOFA, and IntegrAO more consistently recovered significant Reactome, oncogenic, and tissue-relevant gene signatures. In single-cell multiomics benchmarks, method rankings were more dataset dependent, but DIABLO performed consistently well across all case studies, while RGCCA also showed strong performance in specific settings. Computational analyses further showed that DIABLO and MOFA had the most favorable runtime and memory profiles, whereas multiview was the most time-intensive and IntegrAO the most memory-demanding.

**Conclusions:** MESSI provides a reproducible, extensible, and equitable framework for benchmarking multimodal integration methods under a common model assessment strategy. Our results indicate that no single method is uniformly optimal across datasets and objectives; instead, method choice should balance predictive performance, biological interpretability, and computational efficiency. MESSI establishes a foundation for transparent benchmarking and future extensions to broader multimodal learning tasks.

## 1 Background

Technological advances now enable comprehensive profiling of diverse molecular and biomedical modalities, including gene expression, DNA methylation, proteins, metabolites, imaging-derived features, and electrophysiological signals, from the same biological samples. This paradigm, commonly referred to as multimodal data integration, treats each modality as a complementary layer of biological information. Integrating these heterogeneous data sources offers the potential to improve mechanistic understanding of disease and enhance strategies for diagnosis, prognosis, prevention, and therapeutic intervention [1]. Despite its promise, multimodal integration remains methodologically challenging. Each modality differs in scale, distributional properties, dimensionality, noise structure, and missingness patterns, and integration strategies must align with the downstream objective, whether classification, clustering, survival modeling, or biomarker discovery.

Integration in multi-omics and multimodal studies can be conceptualized along two complementary axes [2]: data structure (samples *N* and/or features *P*) and integration approach (**Fig. 1**). From a data-structure perspective, integration strategies are often categorized as horizontal (*P*), vertical (*N*), or mosaic (*NP*) [3–5]. Horizontal integration combines datasets (or views) measured on different samples (*N* differs) but sharing the same or comparable features (*P* is aligned), as in meta-analyses across cohorts or studies profiling the same genes. Vertical integration, in contrast, combines multiple modalities measured on the same set of samples (*N* is shared, *P* differs across modalities), such as transcriptomics, proteomics, and metabolomics collected from the same individuals. This setting enables direct modeling of cross-modality relationships within individuals and is the focus of this study. Mosaic integration addresses partially overlapping designs in which datasets differ in both samples and features, with only subsets shared across modalities or studies, requiring methods that can accommodate missing views and incomplete pairing across *N* and *P*.

**Fig. 1.**
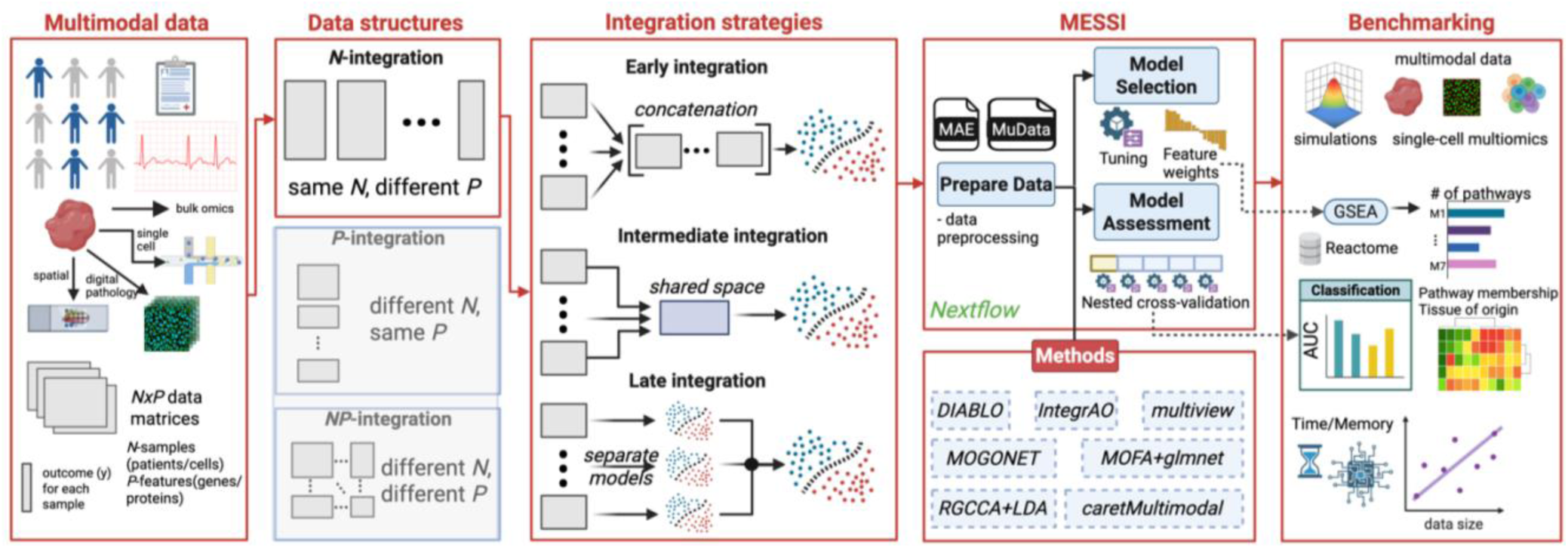
Overview of the MESSI framework for benchmarking data integration methods for outcome prediction. Multimodal biomedical datasets consisting of multiple molecular and clinical modalities measured across shared samples are first organized according to their data structure, including *N*-integration (same samples, different feature sets), *P*-integration (different samples, same features), and *NP*-integration (different samples and different features). These datasets can be integrated using different strategies, including early integration (feature concatenation), intermediate integration (learning a shared latent representation), or late integration (combining predictions from modality-specific models). The MESSI framework implements a reproducible Nextflow-based pipeline that performs data preparation, model selection, and model assessment using nested cross-validation. Multiple multimodal integration methods can be evaluated within this framework (e.g., DIABLO, IntegrAO, multiview cooperative learning, MOGONET, MOFA+glmnet, RGCCA+LDA, and caretMultimodal). Benchmarking outputs include predictive performance, biological interpretability through pathway enrichment analysis, and computational efficiency metrics such as runtime and memory usage across datasets of varying size and modality composition.

Orthogonal to this structural distinction, integration strategies for outcome prediction are categorized according to when modalities are combined within the modeling pipeline: early, intermediate, or late integration [6,7]. Early integration, also known as feature-level integration, concatenates modalities at the input stage after preprocessing and normalization. All features are merged into a single matrix and modeled jointly, allowing direct modeling of cross-modality interactions. However, this approach can substantially increase dimensionality and exacerbate overfitting in the common large *p*, small *n* setting. Modalities with many features may dominate the representation, and improper scaling can introduce imbalance. Intermediate integration, or representation-level integration, combines modalities through learned embeddings or latent variables. Each modality is first modeled separately to extract lower-dimensional representations using projection methods, latent factor models, or neural encoders, which are then aligned or fused into a shared space. This strategy can improve robustness by reducing dimensionality and explicitly modeling shared and modality-specific variation, although it introduces additional modeling complexity and tuning requirements. Finally, late integration trains modality-specific models independently and combines their predictions using ensemble strategies such as averaging or stacking. While this preserves flexibility within each modality, it may fail to capture deeper cross-modal interactions because integration occurs only at the prediction stage.

The diversity of integration paradigms has led to the development of numerous computational frameworks spanning statistical learning and deep learning architectures [8–14] (see **Table 1** for examples). While these methods aim to leverage complementary biological signals across modalities, they differ substantially in assumptions, optimization objectives, supervision strategies, and output formats. Several benchmarking studies [15–17] have compared subsets of integration methods across tasks using public databases such as The Cancer Genome Atlas (TCGA) [18] and Gene Expression Omnibus (GEO) [19]. However, many benchmarks are limited by inconsistent preprocessing, heterogeneous evaluation strategies, restricted method coverage, and lack of reproducibility. Statistical learning approaches are often evaluated using cross-validation, whereas deep learning models frequently rely on fixed train, validation, and test splits. Such inconsistencies complicate interpretation because performance differences may reflect evaluation design rather than inherent methodological strengths.

**Table 1:** List of available multiomics integration methods.

| **Method** | **Type** | **Language** | **Package** | **Reference** |
| --- | --- | --- | --- | --- |
| **BEAM** | Ensemble | R | yes | [21] |
| **caretMultimodal** | Ensemble | R | Yes | [22] |
| **Cooperative Learning** | Penalized regression | R | yes | [23] |
| **DIABLO** | Canonical Correlation Analysis | R | yes | [24] |
| **eipy** | Ensemble | Python | yes | [25] |
| **GOAT** | Deep learning | Python | no | [26] |
| **IntegratedLearner** | Ensemble | R | yes | [27] |
| **IntegrAO** | Deep learning | Python | yes | [28] |
| **JointNMF** | Matrix factorization | R | yes | [29] |
| **MOFA+** | Latent factor model | R/Python | yes | [30] |
| **MOGONET** | Deep learning | Python | no | [31] |
| **mowgli** | Optimal transport | Python | yes | [32] |
| **multiVI** | Deep learning | Python | yes | [33] |
| **RGCCA** | Canonical Correlation Analysis | R | yes | [34] |
| **SLIDE** | Latent factor model | R | yes | [35] |
| **Stabl** | Ensemble | Python | no | [36] |
| **UnitedNet** | Deep learning | Python | no | [37] |

DIABLO (Data Integration Analysis for Biomarker discovery using Latent cOmponents), MOFA+ (Multi-Omics Factor Analysis), RGCCA (Regularized Generalized Canonical Correlation Analysis), IntegrAO (Integrative Analysis with Optimization), IntegratedLearner (Integrated Learner ensemble framework), Stabl (Stability-based Learning), Cooperative Learning (Cooperative penalized regression framework for multi-view data), MOGONET (Multi-Omics Graph cOnvolutional NETwork), GOAT (Graph Omics Attention neTwork), mowgli (Multi-Omics Wasserstein Graph-based Latent Integration), BEAM (Bootstrap-based Ensemble Analysis Method), SLIDE (Sparse Latent Integration for Data Embedding), JointNMF (Joint Non-negative Matrix Factorization), eipy (Ensemble Integration in Python), multiVI (Multi-modal Variational Inference), UnitedNet (Unified Encoder-Decoder Network), and caretMultimodal (Classification And Regression Training Multimodal framework).

Recent efforts have sought to address these limitations through community-driven, reproducible benchmarking infrastructures. For example, the Open Problems in Single-Cell Analysis platform provides a living, extensible framework in which datasets, methods, and evaluation metrics are systematically organized, containerized, and automatically executed within a shared cloud infrastructure [20]. By decoupling evaluation from individual method developers and enabling continual updates as new algorithms emerge, such platforms improve transparency, standardization, and interoperability across studies. However, these frameworks primarily focus on harmonizing tasks, datasets, and metrics rather than enforcing a fully comparable nested cross-validation procedure within each dataset to estimate generalization error. Consequently, while they substantially improve reproducibility of execution, they do not fully resolve the challenge of obtaining unbiased and directly comparable out-of-sample performance estimates across heterogeneous modeling paradigms and hyperparameter tuning strategies.

To address these limitations, we developed *Multimodal Experiments with SyStematic Interrogation* (MESSI), a reproducible Nextflow-based benchmarking framework that enforces a consistent, leakage-free model assessment strategy across diverse classes of multimodal integration methods (**Fig. 1**). Grounded in nested cross-validation, MESSI ensures unbiased hyperparameter tuning and generalization estimates regardless of whether the underlying method follows early, intermediate, or late integration principles. The pipeline standardizes input and output contracts, accommodates heterogeneous hyperparameter grids, and supports bulk molecular data, multimodal biomedical signals, and single-cell multimodal data. By providing a containerized and extensible implementation, MESSI enables transparent, reproducible, and equitable comparisons. In this work, we describe the design and implementation of the framework and apply MESSI to the vertical integration problem, where multiple modalities are captured for the same *N* samples, with a particular emphasis on classification tasks. We benchmark representative statistical and deep learning methods across bulk multi-omics, bulk multimodal, and single-cell multi-omics datasets. Using this unified evaluation hub, we highlight insights into method performance and biological enrichment that emerge only through systematic interrogation.

## 2 Results

### 2.1 MESSI pipeline

The MESSI pipeline (**Fig. 2**) comprises of four main steps: 1) data preparation, 2a) index splitting, 2b) model assessment, 3) model selection and 4) output collection and reporting. MESSI is implemented using the Nextflow [38] domain-specific language, which is widely adopted in bioinformatics for scalable workflow orchestration. Compared to traditional workflow management systems such as Snakemake [39] or GNU Make [40], Nextflow decomposes complex workflows into modular components and connects them via asynchronous channels that govern the flow of the pipeline. This architecture enables each integration method to be encapsulated as an independent workflow with method-specific parameters, while sharing a common evaluation infrastructure. As a result, the pipeline supports not only benchmarking purposes but also broader multi-purpose applications. Additionally, this modular design facilitates rapid, efficient, and reproducible testing and maintenance of codebases.

**Fig. 2:**
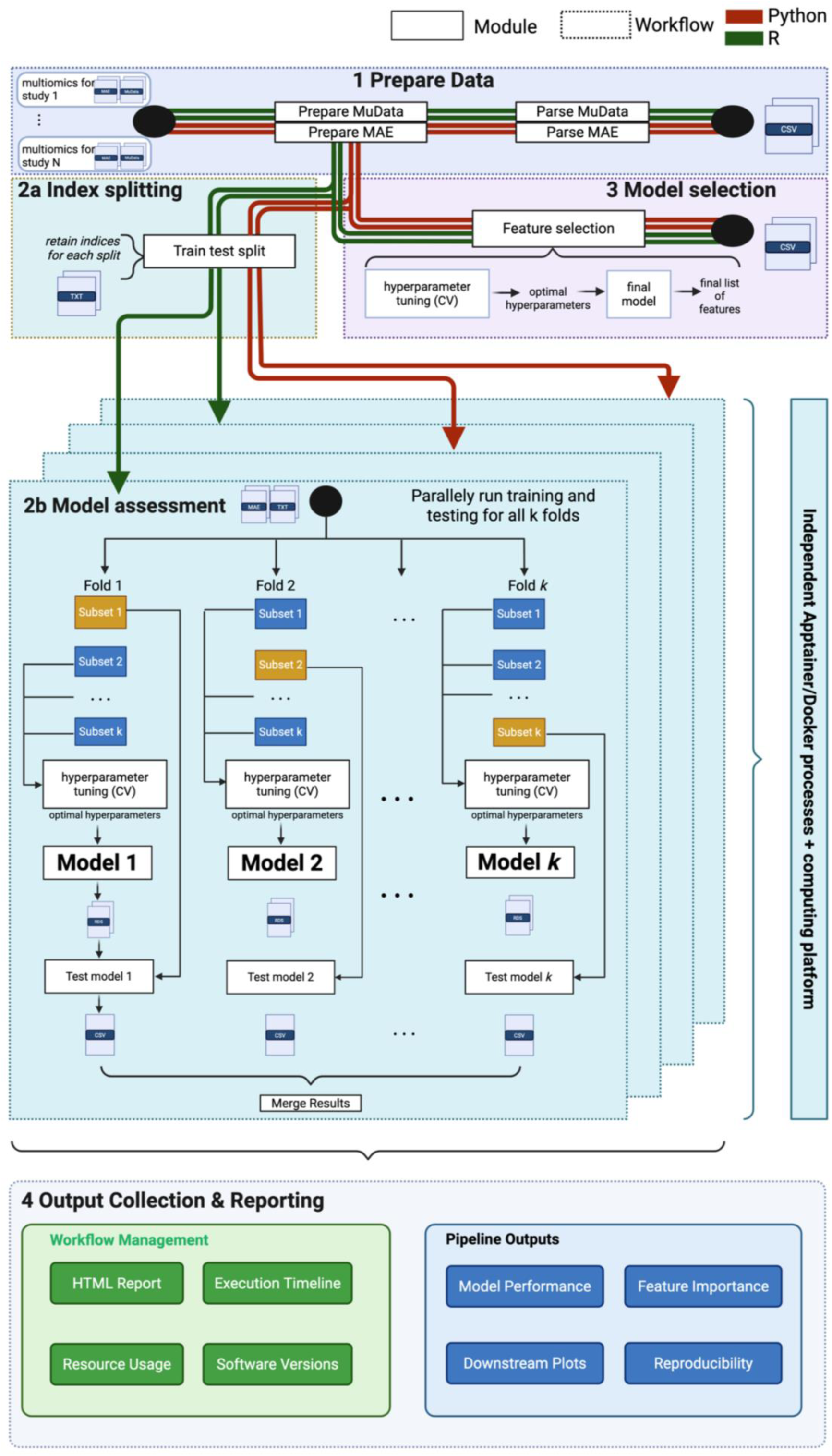
Architecture of the MESSI framework. MESSI employs a modular design composed of four stages: (1) data preparation, (2a) index splitting, (2b) model assessment via *k*-fold cross-validation, (3) model selection and feature extraction, and (4) output collection and reporting. Each stage comprises independent modules (solid boxes) with defined inputs, outputs, and scripts. Workflows (dashed boxes) compose one or more modules and can be nested. Coloured lines denote data flows in R (green) and Python (orange), enabling seamless interoperability across language-specific method implementations. Independent container environments ensure full reproducibility across computing platforms. The parallelizable architecture supports arbitrary combinations of methods, datasets, and folds, scaling efficiently on local machines, HPC clusters, and cloud platforms.

A defining feature of MESSI is its use of containerization technologies such as Docker and Singularity [41], which address common reproducibility challenges. Each time the pipeline is executed, containers are instantiated for every process defined within the workflows. These containers maintain identical operating system configurations and software versions required for the specified computations. Consequently, reproducibility and portability are ensured, as each run is executed within a fully encapsulated and consistent computational environment. Furthermore, Nextflow generates a unique working directory for each process executed within these containers, directing all input/output operations to this designated location without modifying the original input files. This design prevents accidental overwriting of raw data or critical input files.

The independence between processes naturally enables parallelizable computation. This characteristic, combined with Nextflow’s ability to resume interrupted or failed processes, makes debugging or modifying computational settings for individual methods straightforward and modular. Ultimately, this significantly reduces the time and computational burden associated with large datasets or long-running analyses, compared to sequential execution of complex scripts that may fail unpredictably and are difficult to recover.

In terms of interoperability, Nextflow allows the pipeline to run across a wide range of computing platforms, including personal computers, high-performance computing clusters such as SLURM or PBS, and cloud platforms such as AWS and Google Cloud, without requiring modifications to the underlying code logic. This flexibility is achieved through resource and parameter configuration profiles, allowing users to focus on specifying computational resources rather than altering code. An additional important feature is the flexible configuration system, which enables users to run the full collection of methods or select only those of interest for benchmarking. This separation of computational logic from resource configuration ensures that benchmarking results remain portable and reproducible across institutional environments.

Collectively, these features make benchmarking integration methods streamlined and efficient. As data flow through interconnected subworkflows and modules in a factory-like manner, users primarily need to ensure that input data are correctly formatted, after which MESSI manages the remainder of the process. In the following sections, we describe the main components of the pipeline and their implementation, as illustrated in **Fig. 2**.

#### 2.1.1 Prepare data

The first stage standardises heterogeneous input datasets into common analytical formats. Integration methods differ widely in their expected input formats, necessitating a unified preprocessing layer. MESSI accepts input as a CSV manifest specifying dataset identifiers and paths to compressed archives, each containing data in both MultiAssayExperiment (MAE) [42] format for R-based methods and MuData [43] format for Python-based-methods:

The input of the pipeline is a csv like the following:

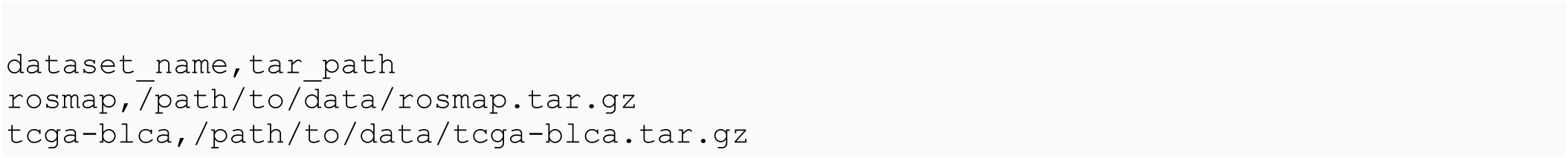

This csv input allows user to specify multiples datasets as long as it follows the requirement of providing an identifier, and a path to a compressed tar which consists of two key file formats .h5 and .h5mu. An example of a dataset compressed tar would be:

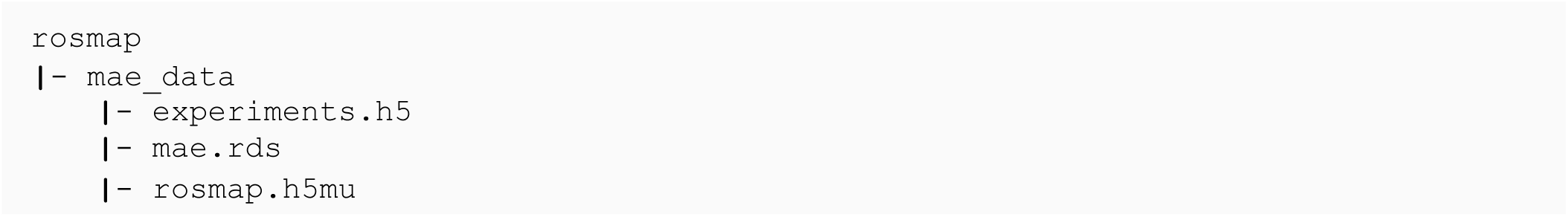

Upon ingestion, the pipeline performs the following steps: (1) decompression of each dataset archive into an isolated working directory; (2) parallel preprocessing of both MAE and MuData representations, including removal of missing observations, variance-based feature filtering, exclusion of near-zero variance features, and conversion of response variables to binary encoding where appropriate; and (3) extraction of key dataset metadata, including omics modality names, feature dimensions, sample counts, and class proportions. Both the R and Python preprocessing pathways are rigorously validated for numerical consistency, ensuring that downstream methods receive identical data regardless of the implementation language. Through these two core APIs, multimodal datasets can be seamlessly transformed between R and Python environments without loss of information or structural integrity. This represents an important step toward standardizing file formats for multimodal data, as current studies often rely on heterogeneous and incompatible data structures that hinder reproducibility. By supporting interoperable MAE and MuData representations, our framework promotes a unified and reproducible data standard. Furthermore, leveraging these two ecosystems facilitates the adoption of integration methods implemented in different programming languages, enabling users to work within their preferred computational environment while maintaining compatibility across tools.

#### 2.1.2 Index splitting

To support different types of analyses, we implement data splitting using a standard cross-validation strategy based on the MuData representation. This choice leverages the robust support available in Python libraries, particularly scikit-learn, which provide well-tested APIs such as StratifiedKFold for reproducible and class-balanced partitioning. Using this framework, we generate folds under a default 5-fold cross-validation setting and record the test-set indices for each fold as `.txt` files for downstream use. The number of folds can be adjusted through the Nextflow configuration parameter `params.k`. This externalized splitting strategy ensures that all methods operate on identical train–test partitions, eliminating confounding variation arising from method-specific splitting procedures or discrepancies in random seed handling. By persisting fold indices as text files, we guarantee transparency and reproducibility across runs and implementations.

Furthermore, the design naturally supports nested cross-validation. An outer loop evaluates generalization performance, while an optional inner loop implemented within each method optimizes hyperparameters. The inner cross-validation procedure is controlled via the `params.inner_cv` configuration parameter, enabling flexible benchmarking under both simple and nested cross-validation regimes.

#### 2.1.4 Model selection

This workflow implements model selection, the process of choosing among candidate models or tuning parameter configurations based on their estimated predictive performance [44]. In our framework, model selection therefore includes identifying the optimal model specification (e.g. hyperparameters) as well as the associated feature weights. When sparsity-inducing methods are used, this step additionally determines which features are retained (non-zero coefficients) and which are excluded. The model selection workflow operates directly on the fully preprocessed data generated during the “Prepare Data” stage, as it leverages the complete dataset to optimize tuning parameters and estimate final feature effects. This workflow can be executed either prior to or concurrently with the model assessment stage. Similar to the model assessment component, it consists of independent method-specific subworkflows, enabling parallel execution for scalability and computational efficiency. For each dataset–method combination, the output is a structured table containing the selected features along with their corresponding coefficients, loadings, or importance scores, depending on the modeling framework. In sparse models, this table explicitly reflects the subset of features assigned non-zero weights. These results are aggregated across all evaluated methods using the full dataset (without train–test partitioning) and subsequently forwarded to the reporting stage for downstream summarization, comparison, and visualization.

#### 2.1.5 Model assessment

The model assessment stage constitutes the computational core of the pipeline and implements nested cross-validation, estimating the generalization error of a model, that is, it’s expected predictive performance on new, unseen data [44]. Each integration method is implemented as an independent workflow comprising three required modules: preprocess, train, and predict. While additional modules may be included depending on the method, these three components are mandatory. The input to each method consists of either the MAE or MuData representation of the original dataset, depending on the programming language in which the method is implemented. Treating each method as a self-contained workflow provides flexibility to define method-specific parameters within the workflow itself, while also allowing global control via Nextflow configuration files. This modular structure further enables extension of individual methods, for example, by incorporating additional modules for exploratory data analysis, visualization, or other method-specific utilities. Within the cross-validation stage, the preprocess module converts the standardized MAE/MuData objects generated in the “Prepare Data” stage into method-specific input formats. This step is essential because different methods require distinct data structures (e.g., matrices, tensors, lists, or custom objects). After preprocessing, data are partitioned according to the test-set indices generated during the “Index Splitting” stage. This produces partitions of the form data*_i_*–fold*_j_*, where *i* = 1, …, *N* indexes datasets and *j* = 1, …, *K* indexes folds (with *K* = 5 by default). In the train module, the training subset of data*_i_*–fold*_j_* is used to fit the model. By default, methods are trained using their standard parameter settings unless nested cross-validation is enabled. When nested cross-validation is activated, an inner cross-validation loop is performed within the training set to conduct model selection (e.g., hyperparameter tuning). The optimal configuration identified in the inner loop is then used to retrain the model on the full outer training folds. The held-out outer test fold is subsequently passed to the predict module, which generates predictions using the trained model. This nested structure ensures that hyperparameter tuning does not leak information from the outer test fold, thereby providing an unbiased estimate of generalization performance. All method–dataset–fold combinations (*M* × *N* × *K*, where *M* is the number of methods, *N* the number of datasets, and *K* the number of folds) are executed in parallel when computational resources permit. Each method produces standardized outputs comprising predicted probabilities, sample identifiers, method names, and dataset identifiers, thereby enabling uniform downstream evaluation. Performance metrics are aggregated per method across all datasets and folds and subsequently forwarded to the reporting stage.

#### 2.1.5 Configuration of parameters

MESSI’s configuration system leverages Nextflow’s profile mechanism, enabling hierarchical composition of parameter sets for different computing environments, containerisation strategies, and dataset configurations: Each profile may inherit from others and override parameter values, supporting a hierarchical and modular approach to configuration. In our implementation, important options such as skipping specific methods, adjusting the number of cross-validation folds, modifying method-specific hyperparameters, or adapting to site/platform-specific computing environments are all controlled through Nextflow profiles. Each profile is defined in a `.config` file using a Groovy-based syntax similar to YAML, but with the additional ability to evaluate Nextflow expressions dynamically. For example:

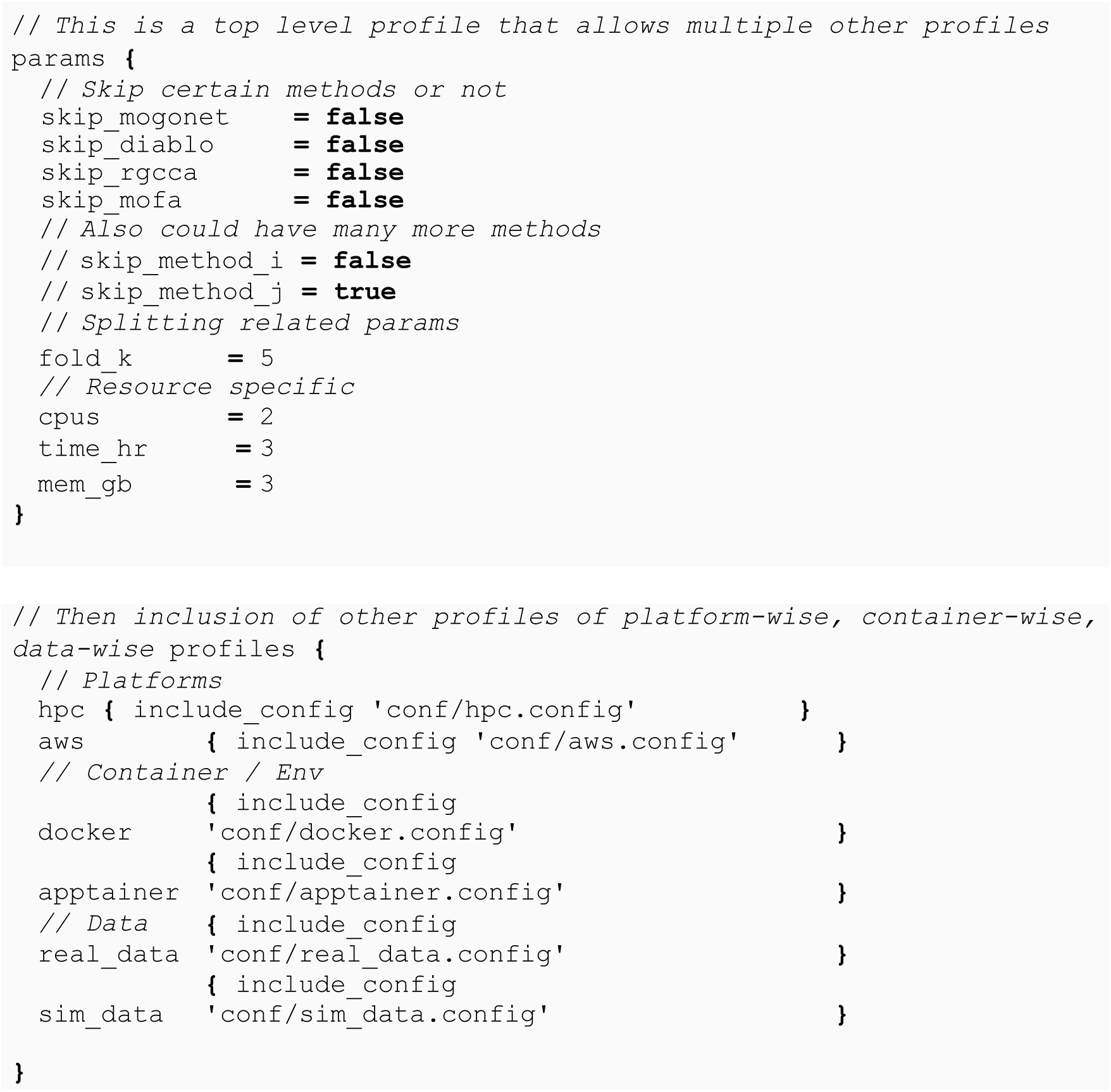

Execution can then be customized by chaining multiple profiles to create tailored configurations. For instance:

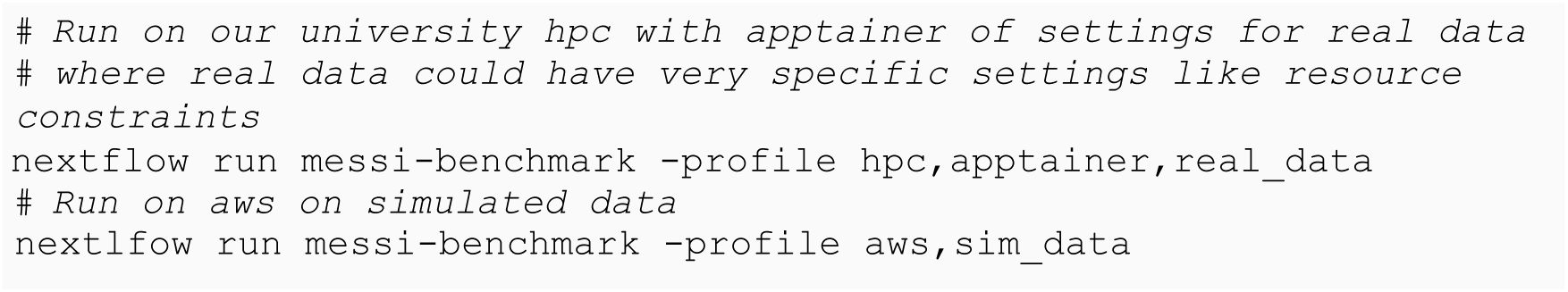

This modular and declarative approach enables extensive flexibility for users, allowing parameter tuning, environment adaptation, and reproducible configuration from a single control script. Changing a single parameter in a profile automatically propagates to all pipeline components where that parameter is used. Furthermore, this structure facilitates systematic exploration of hyperparameters and preserves a complete record of configurations used in each run, critical for reproducibility, benchmarking, and downstream analysis.

#### 2.1.6 Integration methods

MESSI currently includes a set of representative integrative methods spanning intermediate and late integration strategies. Given the broad scope of multimodal data integration, we focused this benchmark on vertical integration (N-integration) for binary classification tasks aimed at disease or condition outcome prediction. The intermediate integration methods included Cooperative Learning (Multiview), Data Integration Analysis for Biomarker discovery using Latent cOmponents (DIABLO), Integrative Analysis of Omics data (IntegrAO), Multi-Omics Factor Analysis (MOFA), and Regularized Generalized Canonical Correlation Analysis (RGCCA), while the late integration methods included an ensemble of penalized regression models implemented with glmnet via caretMultimodal and Multi-Omics Graph cOnvolutional NETworks (MOGONET). Although intermediate integration methods are generally designed to extract information shared across views, some methods, including DIABLO, Multiview, and RGCCA, allow the user to control the extent to which shared structure is emphasized. In DIABLO and RGCCA, this is specified through a design matrix that defines the degree of association between views. For benchmarking, we evaluated the two extremes: no association between views (null design) and strong association between views (full design), denoted as DIABLO-N, DIABLO-F, RGCCA-N, and RGCCA-F. For Multiview, the corresponding correlation parameter was tuned during model selection. MOFA and RGCCA are unsupervised methods, so their extracted latent factors were subsequently used for binary classification with either ridge regression (MOFA+glmnet) or linear discriminant analysis (RGCCA+LDA). For latent factor-based methods such as MOFA, RGCCA, and DIABLO, we retained two components rather than tuning the number of components, to enable a fairer comparison across methods. We used the notation DIABLO-N1, DIABLO-F1, RGCCA-N1+LDA, RGCCA-F1+LDA, and MOFA-OneK1K cohort dataset1+glmnet, and similarly DIABLO-N2, DIABLO-F2, RGCCA-N2+LDA, RGCCA-F2+LDA, and MOFA-2+glmnet, to refer to the first and second components, respectively. For classification, both components were used jointly, hence the simplified notation DIABLO-N, DIABLO-F, RGCCA-N+LDA, and RGCCA-F+LDA; however, for feature selection and gene set enrichment analysis, component-specific notation was retained because feature weights were analyzed separately for each component. Finally, because some methods inherently perform variable selection whereas others do not, we did not tune sparsity parameters across methods. Instead, we used feature weights as a common basis for comparison and downstream gene set enrichment analysis.

### 2.2 Simulation studies

To establish ground truth for method evaluation in a binary classification setting, we conducted simulation studies with known data-generating processes (see Methods), enabling precise assessment of both predictive performance and feature recovery. We generated multiomics datasets comprising three omics views, each containing *p*=3000 features measured on *n*=1,000 samples, with 10% of features designated as true signal variables (based on one underlying latent factor) associated with a binary outcome. The outcome labels were generated from a model in which only the signal features contributed to class separation. Signal strength and inter-omics correlation were varied systematically to create a factorial experimental design spanning regimes of no signal, low signal, and high signal, crossed with low, medium, and high inter-omics correlation.

To assess how different multimodal integration methods perform in simulation studies, we evaluated both their classification performance and their ability to recover true signal variables. For classification performance, we quantified how well each method identified the correct response class using the area under the receiver operating characteristic curve (AUC). For feature selection performance, we evaluated how frequently top-ranked variables corresponded to the true signal variables specified in the simulation design, using metrics analogous to sensitivity and specificity by assessing whether highly ranked features belonged to the predefined signal set versus noise variables. To account for randomness and variability in model training and data splitting, we implemented 5-fold cross-validation. Full details of the simulation framework, including data generation procedures and model implementation, are provided in an accompanying R package, with additional technical details described in the Methods section.

The simulation results revealed three main findings. First, all methods were well calibrated in the absence of a discriminative signal, with mean AUC values centred around 0.50 across methods, indicating no systematic tendency toward false positive classification and supporting the validity of downstream comparisons under informative settings (**Fig. 3A**). Second, performance diverged under low-signal conditions, which are likely the most representative of real biological datasets where effect sizes are modest. Under low signal, all methods achieved strong discrimination, with the exception of MOFA-1+glmnet and RGCCA-N1+LDA, which performed less well. This pattern suggests that unsupervised or weakly supervised approaches were less effective at recovering modest class-discriminative signal. In particular, MOFA may prioritize dominant sources of shared variation rather than outcome-related structure, while RGCCA appears sensitive to the design specification used to define inter-omic relationships. The stronger performance of RGCCA-F1+LDA relative to RGCCA-N1+LDA suggests that the full design matrix partly rescued classification by better capturing shared structure across omics blocks and passing more informative latent components to the downstream LDA classifier. Under high-signal conditions, all methods converged to near-perfect classification performance, as expected when class structure is strong and unambiguous. Inter-omics correlation had little effect on AUC across signal settings, indicating that classification performance was broadly robust to the degree of shared structure between modalities. Feature-selection analyses further highlighted important differences between methods (**Fig. 3B**). For true variables, as signal strength increased, DIABLO variants, RGCCA variants, IntegrAO, and caretMultimodal showed high sensitivity for recovering true variables, whereas Multiview was more moderate and MOFA-1+glmnet and MOGONET recovered fewer true signals, particularly under low signal. However, the noise-variable panel indicates that false positive feature selection remained substantial for several methods, such that strong recovery of true variables did not always correspond to strong exclusion of uninformative features. Runtime comparisons showed that Multiview and IntegrAO were the most computationally intensive during model selection, each requiring more than 1,000 seconds, whereas MOFA, DIABLO, and MOGONET were among the fastest methods (**Fig. 3C**). Notably, model assessment required a wall-clock time similar to model selection rather than substantially longer. This reflects the parallelized design of the MESSI workflows, in which computations for different folds are executed simultaneously, making total runtime roughly comparable to that of a single model-selection run.

**Fig. 3.**
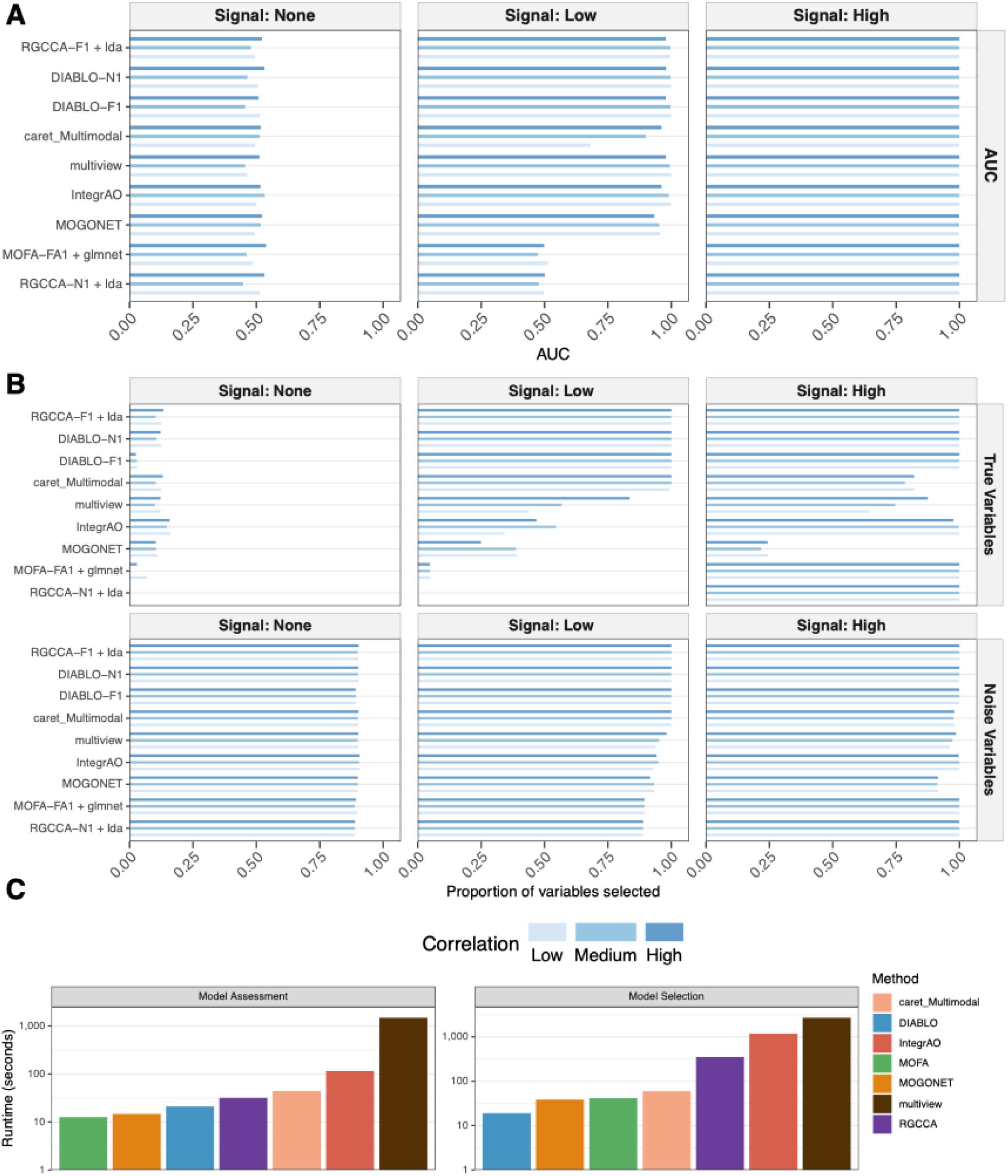
Benchmarking multimodal integration methods on simulated datasets across varying signal strengths. A. Classification performance, measured as the area under the receiver operating characteristic curve (AUC) from 5-fold cross-validation, for each method under no-signal, low-signal, and high-signal simulation settings. B. Feature-selection behavior, shown as the proportion of true variables and noise variables selected across simulation settings. C. Computational runtime for the model assessment and model selection stages. Bar shading denotes the correlation structure of the simulated data (low, medium, and high).

### 2.3 Bulk multimodal benchmarking

We benchmarked MESSI across 16 bulk multimodal datasets spanning a wide range of diseases, outcome definitions, sample sizes, and modality combinations (**Fig. 4A**). These datasets included neurodevelopmental, neurodegenerative, cancer, renal, infectious, and metastatic disease settings, with cohort sizes ranging from 7 patients with 149 imaging regions of interest to 599 samples, and positive-class proportions ranging from 0.22 to 0.762 (**Table 2**). Most datasets integrated molecular layers such as mRNA and CpG methylation, often alongside miRNA, protein, copy number, or chromatin-compartment features, while others incorporated non-molecular modalities including imaging, electrical measurements, cellular features, and clinical variables. Feature dimensionality also varied substantially across studies, from fewer than 10 clinical variables to over 19,000 transcriptomic or genomic features. Several datasets contrasted early versus advanced disease stage, whereas others focused on control versus disease, hospitalization status, or primary versus metastatic disease. Collectively, this benchmark captures the diversity and complexity of real-world multimodal biomedical data and provides a robust evaluation of MESSI across small and large cohorts, balanced and imbalanced class distributions, and a broad spectrum of molecular and non-molecular data types.

**Figure 4.**
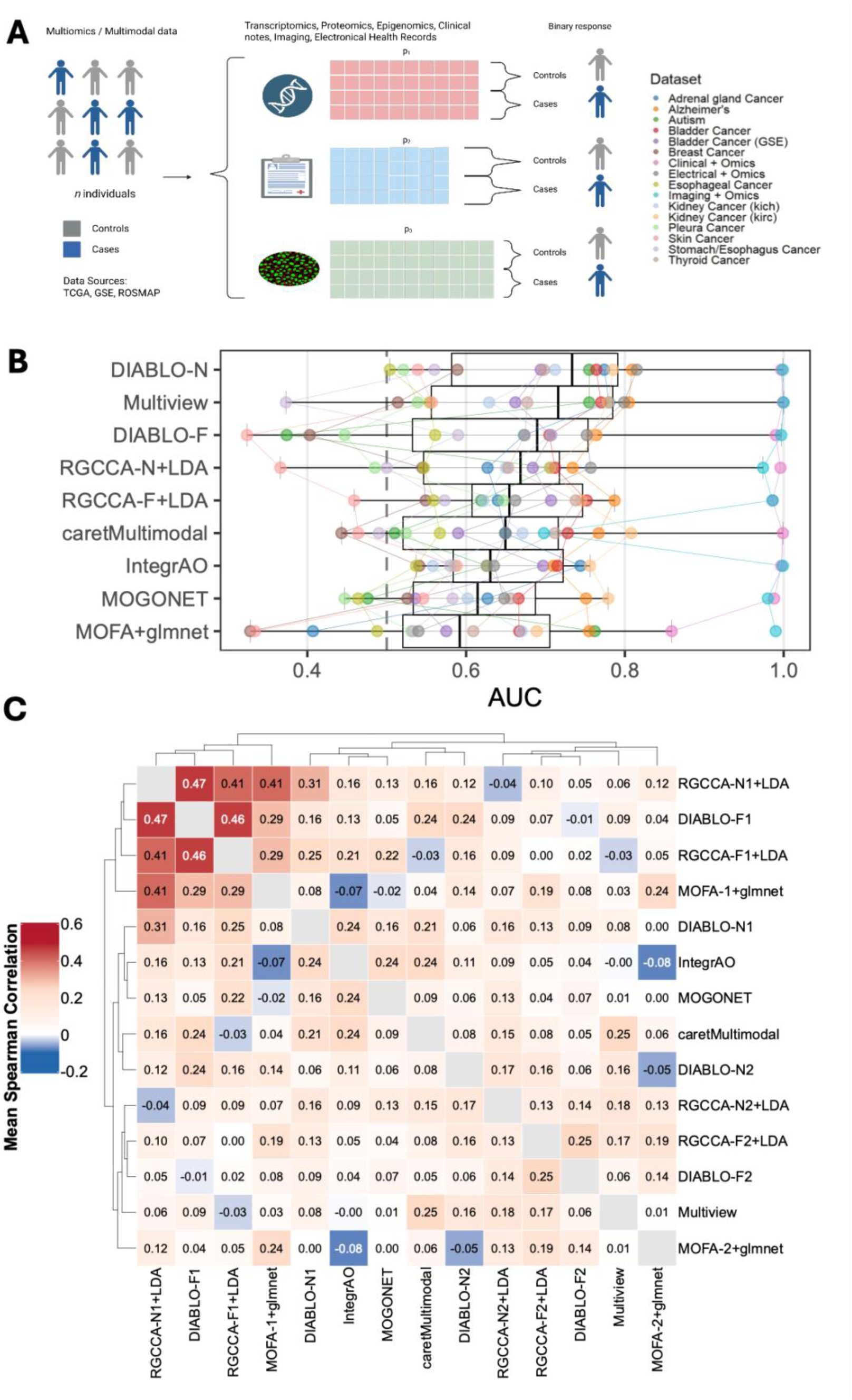
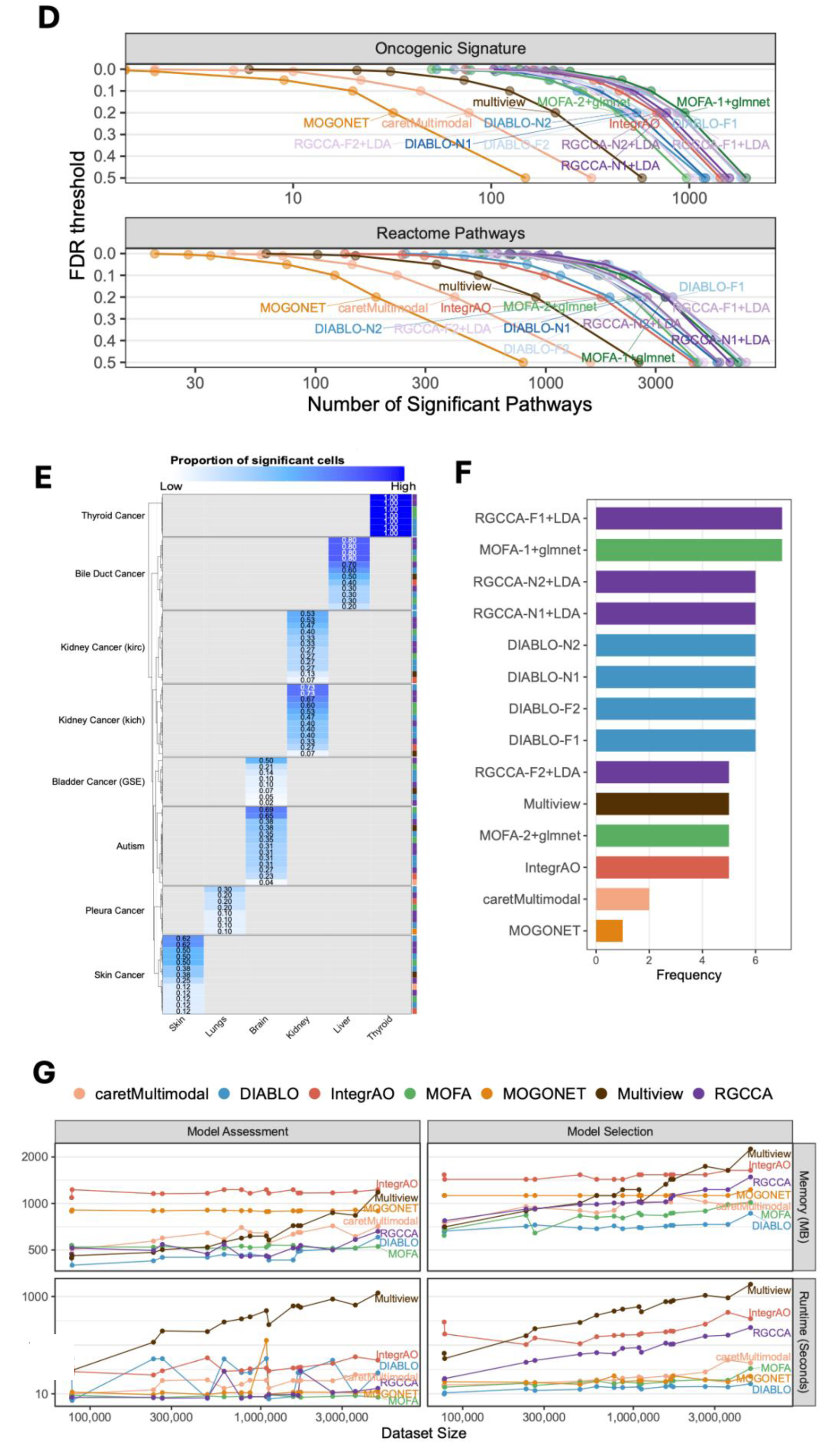
Benchmarking integration methods on bulk multimodal datasets. A. Overview of the bulk multimodal benchmarking design across 16 datasets spanning diverse diseases, outcome definitions, and modality combinations, including molecular and non-molecular data types. B. Classification performance across datasets, shown as AUC for each integration method. C. Heatmap of mean Spearman correlations between feature weights, showing concordance in feature prioritization across methods and latent components. D. Biological enrichment analysis summarizing the number of significant oncogenic-signature and Reactome pathways recovered from ranked feature weights across FDR thresholds. E. Heatmap showing enrichment of method-derived feature rankings for tissue-relevant PanglaoDB cell-marker gene sets across cancer datasets. F. Summary of tissue-of-origin matched associations identified by each method. G. Computational scalability of each method, shown by peak memory usage and runtime during model assessment and model selection as dataset size increases.

**Table 2:**
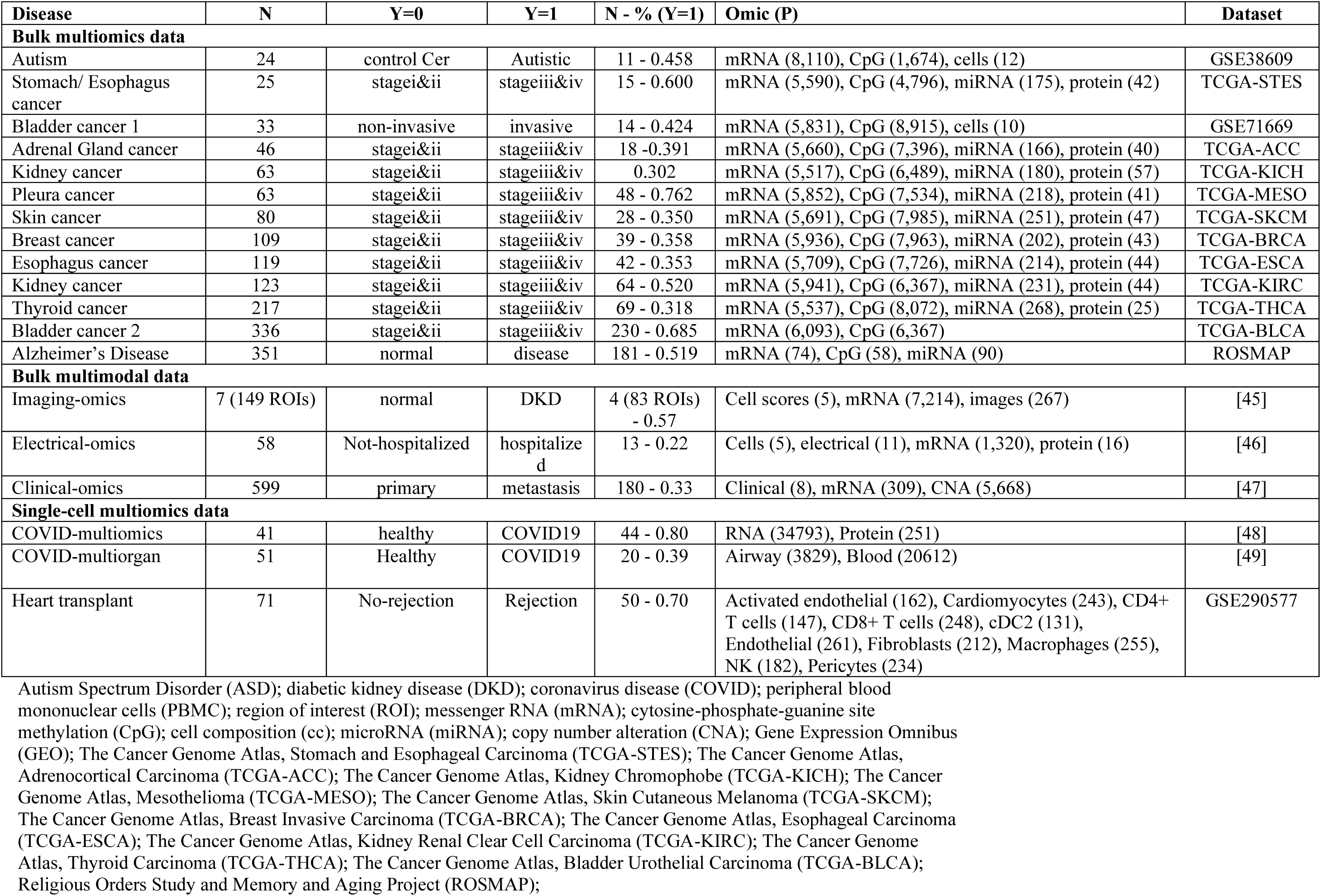
Overview of real datasets to benchmark.

| Disease | N | Y=0 | Y=1 | N - % (Y=1) | Omic (P) | Dataset |
| --- | --- | --- | --- | --- | --- | --- |
| **Bulk multiomics data** |  |  |  |  |  |  |
| Autism | 24 | control Cer | Autistic | 11 - 0.458 | mRNA (8,110), CpG (1,674), cells (12) | GSE38609 |
| Stomach/ Esophagus cancer | 25 | stagei&ii | stageiii&iv | 15 - 0.600 | mRNA (5,590), CpG (4,796), miRNA (175), protein (42) | TCGA-STES |
| Bladder cancer 1 | 33 | non-invasive | invasive | 14 - 0.424 | mRNA (5,831), CpG (8,915), cells (10) | GSE71669 |
| Adrenal Gland cancer | 46 | stagei&ii | stageiii&iv | 18 - 0.391 | mRNA (5,660), CpG (7,396), miRNA (166), protein (40) | TCGA-ACC |
| Kidney cancer | 63 | stagei&ii | stageiii&iv | 0.302 | mRNA (5,517), CpG (6,489), miRNA (180), protein (57) | TCGA-KICH |
| Pleura cancer | 63 | stagei&ii | stageiii&iv | 48 - 0.762 | mRNA (5,852), CpG (7,534), miRNA (218), protein (41) | TCGA-MESO |
| Skin cancer | 80 | stagei&ii | stageiii&iv | 28 - 0.350 | mRNA (5,691), CpG (7,985), miRNA (251), protein (47) | TCGA-SKCM |
| Breast cancer | 109 | stagei&ii | stageiii&iv | 39 - 0.358 | mRNA (5,936), CpG (7,963), miRNA (202), protein (43) | TCGA-BRCA |
| Esophagus cancer | 119 | stagei&ii | stageiii&iv | 42 - 0.353 | mRNA (5,709), CpG (7,726), miRNA (214), protein (44) | TCGA-ESCA |
| Kidney cancer | 123 | stagei&ii | stageiii&iv | 64 - 0.520 | mRNA (5,941), CpG (6,367), miRNA (231), protein (44) | TCGA-KIRC |
| Thyroid cancer | 217 | stagei&ii | stageiii&iv | 69 - 0.318 | mRNA (5,537), CpG (8,072), miRNA (268), protein (25) | TCGA-THCA |
| Bladder cancer 2 | 336 | stagei&ii | stageiii&iv | 230 - 0.685 | mRNA (6,093), CpG (6,367) | TCGA-BLCA |
| Alzheimer's Disease | 351 | normal | disease | 181 - 0.519 | mRNA (74), CpG (58), miRNA (90) | ROSMAP |
| **Bulk multimodal data** |  |  |  |  |  |  |
| Imaging-omics | 7 (149 ROIs) | normal | DKD | 4 (83 ROIs) - 0.57 | Cell scores (5), mRNA (7,214), images (267) | [45] |
| Electrical-omics | 58 | Not-hospitalized | hospitalized | 13 - 0.22 | Cells (5), electrical (11), mRNA (1,320), protein (16) | [46] |
| Clinical-omics | 599 | primary | metastasis | 180 - 0.33 | Clinical (8), mRNA (309), CNA (5,668) | [47] |
| **Single-cell multiomics data** |  |  |  |  |  |  |
| COVID-multiomics | 41 | healthy | COVID19 | 44 - 0.80 | RNA (34793), Protein (251) | [48] |
| COVID-multiorgan | 51 | Healthy | COVID19 | 20 - 0.39 | Airway (3829), Blood (20612) | [49] |
| Heart transplant | 71 | No-rejection | Rejection | 50 - 0.70 | Activated endothelial (162), Cardiomyocytes (243), CD4+ T cells (147), CD8+ T cells (248), cDC2 (131), Endothelial (261), Fibroblasts (212), Macrophages (255), NK (182), Pericytes (234) | GSE290577 |

Autism Spectrum Disorder (ASD); diabetic kidney disease (DKD); coronavirus disease (COVID); peripheral blood mononuclear cells (PBMC); region of interest (ROI); messenger RNA (mRNA); cytosine-phosphate-guanine site methylation (CpG); cell composition (cc); microRNA (miRNA); copy number alteration (CNA); Gene Expression Omnibus (GEO); The Cancer Genome Atlas, Stomach and Esophageal Carcinoma (TCGA-STES); The Cancer Genome Atlas, Adrenocortical Carcinoma (TCGA-ACC); The Cancer Genome Atlas, Kidney Chromophobe (TCGA-KICH); The Cancer Genome Atlas, Mesothelioma (TCGA-MESO); The Cancer Genome Atlas, Skin Cutaneous Melanoma (TCGA-SKCM); The Cancer Genome Atlas, Breast Invasive Carcinoma (TCGA-BRCA); The Cancer Genome Atlas, Esophageal Carcinoma (TCGA-ESCA); The Cancer Genome Atlas, Kidney Renal Clear Cell Carcinoma (TCGA-KIRC); The Cancer Genome Atlas, Thyroid Carcinoma (TCGA-THCA); The Cancer Genome Atlas, Bladder Urothelial Carcinoma (TCGA-BLCA); Religious Orders Study and Memory and Aging Project (ROSMAP);

#### 2.3.1 Bulk multimodal classification performance

We benchmarked the integration methods across bulk multimodal datasets comprising combinations of transcriptomic, proteomic, epigenomic, clinical, and imaging-derived features (**Fig. 4A**). Overall, DIABLO-N and Multiview showed the strongest classification performance, whereas MOFA+glmnet and MOGONET showed the weakest performance (**Fig. 4B**). The difference between the best- and worst-performing methods was >10% in AUC, but given the spread of AUC values across datasets, these differences were not statistically significant between methods. This indicates that although some methods tended to rank higher than others, overall predictive performance was broadly comparable across the methods.

Feature-weight concordance analysis provided additional insight into how methods behaved in the classification setting. Spearman correlations between the feature weights (absolute value), averaged across datasets, showed clustering of related intermediate integration methods, particularly RGCCA, DIABLO, and MOFA (**Fig. 4C**). Notably, component 1 from these methods tended to cluster together, and component 2 also clustered together, whereas the distinction between null and full design matrices had less influence on the clustering. This suggests that the dominant source of similarity across these approaches was the latent component captured rather than the specific correlation design imposed.

#### 2.3.2 Biological enrichment of bulk multimodal features

To assess biological relevance, we performed gene set enrichment analysis (GSEA) using the feature weights from each method–dataset combination and then summarized the number of significant pathways across a range of FDR thresholds (**Fig. 4D**). DIABLO, RGCCA, MOFA, and IntegrAO identified the largest numbers of significant pathways in both the Reactome and oncogenic signature collections, indicating that these methods more consistently recovered biologically coherent signal from the ranked features. In contrast, methods such as MOGONET and caretMultimodal tended to identify fewer enriched pathways. Together, these results suggest that strong biological enrichment was not restricted to the top classifiers and that some methods may be especially useful when interpretability is a primary goal.

We further assessed tissue specificity using PanglaoDB-derived cell-marker gene sets by asking whether method-specific feature rankings were enriched for cell types relevant to the tissue of origin of each cancer dataset (**Fig. 4E**). For example, for skin cancer datasets, we evaluated whether the ranked features were enriched for skin-related cell markers. This analysis provided an orthogonal measure of biological plausibility by linking selected features to expected tissue-specific cellular contexts. Complementing this result, the tissue-of-origin association summary showed that RGCCA, MOFA, and DIABLO identified the greatest number of tissue-matched associations, whereas MOGONET, caretMultimodal, and IntegrAO identified the fewest (**Fig. 4F**). These findings indicate that RGCCA, MOFA, and DIABLO more consistently recovered biologically meaningful, tissue-relevant signals across the benchmark datasets.

#### 2.3.3 Bulk multimodal computational scalability

To evaluate practical feasibility, we examined runtime and peak memory usage during both model assessment and model selection as dataset size increased (**Fig. 4G**). DIABLO and MOFA showed the most favorable computational profiles, with relatively modest increases in runtime and memory usage as dataset size grew. In contrast, Multiview had the highest memory demands and longest runtimes, both of which increased with dataset size.

### 2.4 Single-cell multiomics benchmarking

We benchmarked MESSI across three single-cell multiomics datasets spanning infectious and transplant-related disease settings, with distinct outcome definitions, cohort sizes, and modality structures (**Fig. 5A** and **Table 2**). These included two COVID-19 datasets comparing healthy controls with COVID-19 cases, and one heart transplant dataset contrasting no rejection versus rejection. Cohort sizes ranged from 41 to 71 samples, and positive-class proportions ranged from 0.39 to 0.80. The datasets also differed substantially in feature dimensionality and modality composition. The COVID-multiomics dataset integrated two modalities, including pseudobulk mRNA profiles, and protein profiles. The COVID-multiorgan dataset also comprised two modalities, but at substantially larger scale, including airway mRNA profiles, and PBMC mRNA profiles. In contrast, in the heart transplant dataset, each cell type was treated as a separate modality, resulting in 10 cell types across 71 samples. Collectively, these datasets capture the diversity of single-cell multiomics benchmarking scenarios, spanning moderate to high class imbalance, two- and ten-modality designs, and feature spaces ranging from a few thousand to hundreds of thousands of variables per modality.

**Fig. 5:**
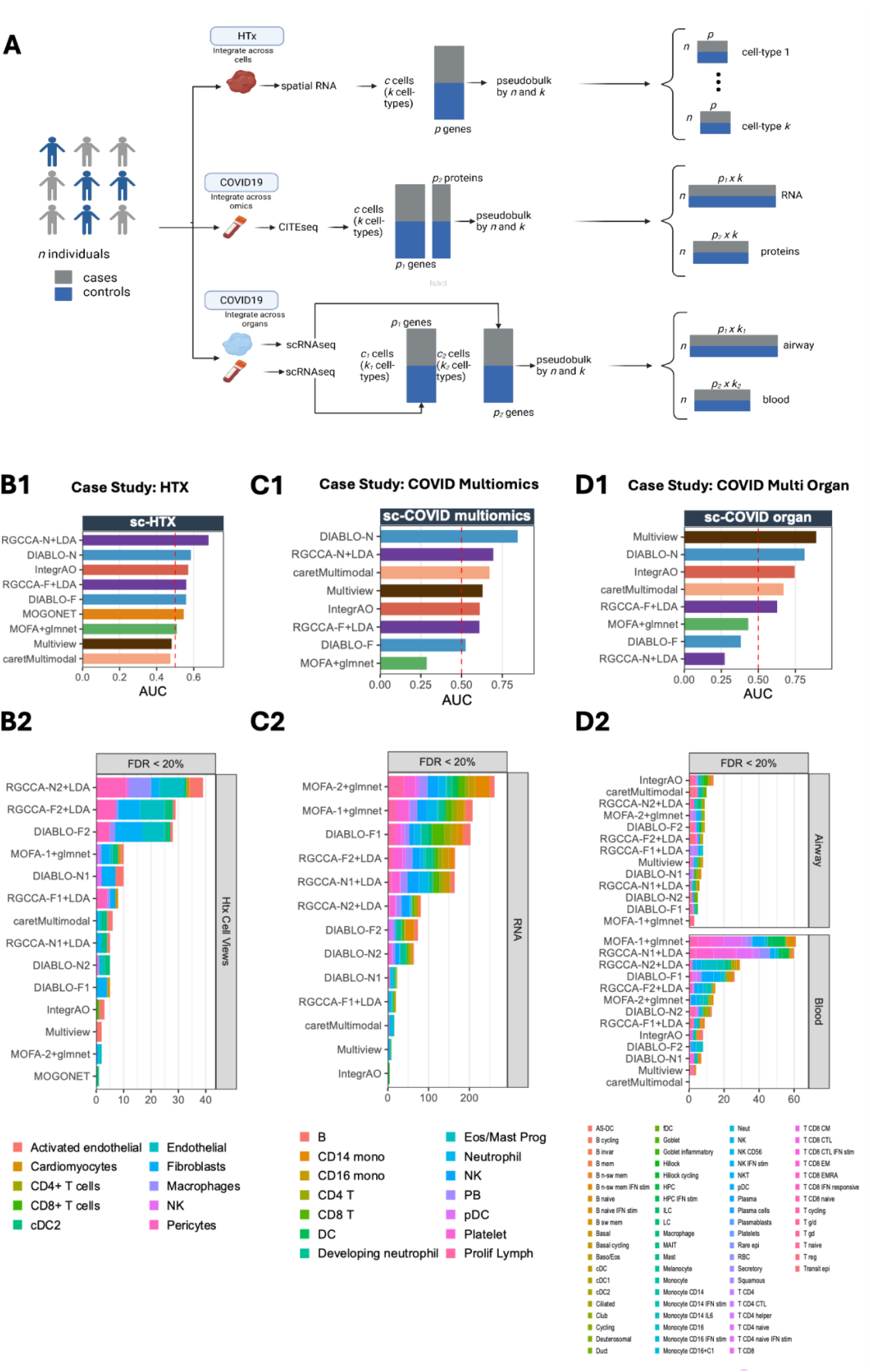
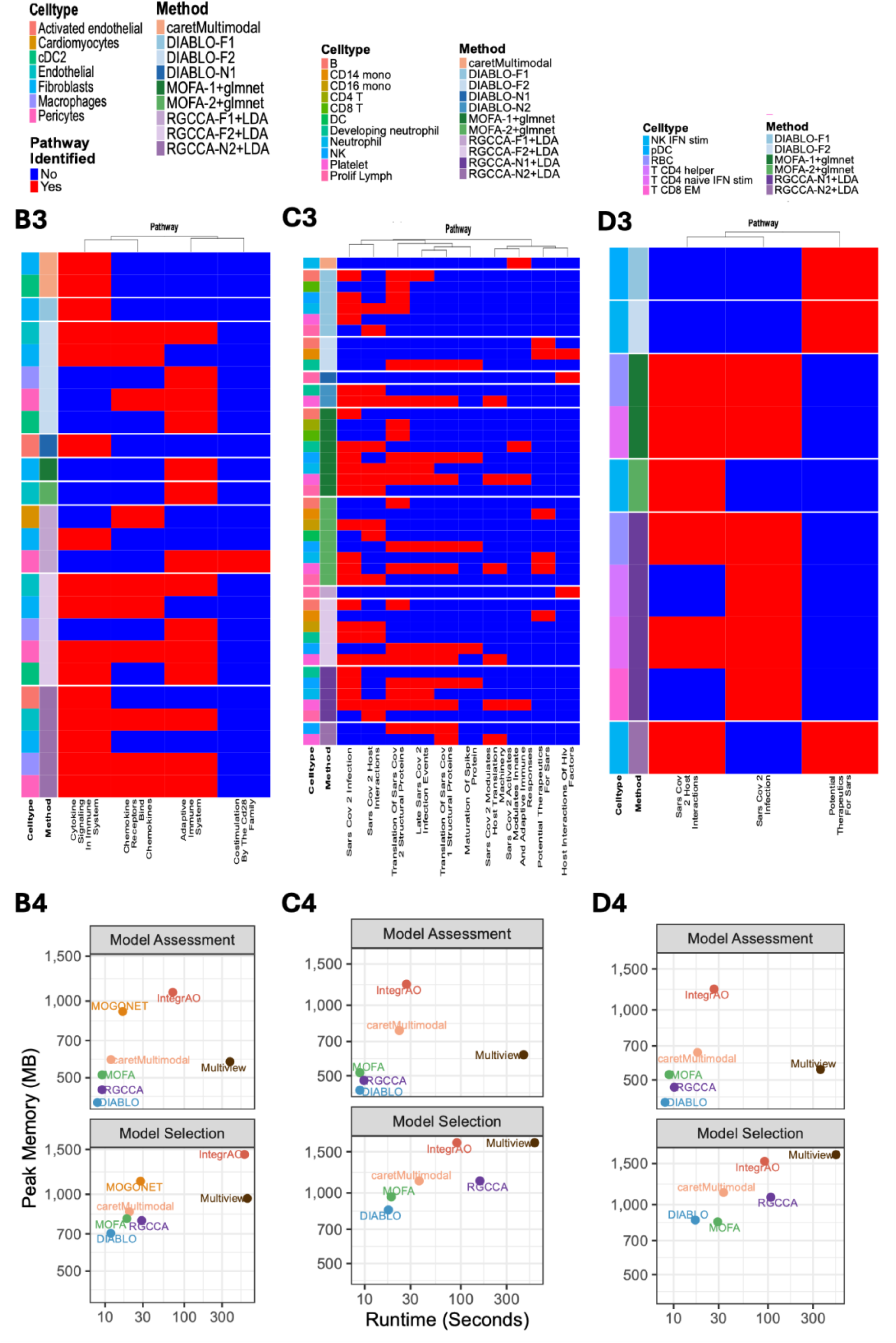
Benchmarking multimodal integration methods on single-cell datasets. **(A)** Overview of the single-cell benchmarking datasets and pseudobulk aggregation strategy. The heart transplant (HTX) dataset comprises spatial transcriptomics data integrated across cell types, the COVID-19 multiomics dataset comprises CITE-seq data (RNA and protein) integrated across omics, and the COVID-19 organ dataset comprises scRNA-seq data from airway and blood integrated across organs. For each dataset, single-cell data were aggregated into pseudobulk profiles by patient and cell type, yielding patient-by-feature matrices for downstream classification of cases versus controls. **(B)** Classification performance AUC from 5-fold cross-validation across integration methods for the three single-cell datasets: sc-HTX (top), sc-COVID multiomics (middle), and sc-COVID organ (bottom). The red dashed line indicates chance-level threshold (AUC = 0.5). Methods are ranked by descending mean AUC within each dataset. **(C)**, Heatmap showing the number of significant pathways (FDR < 0.2) identified per method and cell type in the sc-HTX dataset. Columns represent cell types and rows represent integration methods, with colour intensity reflecting pathway count. **(D)**, Number of significant pathways identified per method in the sc-COVID multiomics dataset, stratified by data modality (protein, left; RNA, right). Stacked bars are coloured by cell type. **(E)**, Number of significant pathways identified per method in the sc-COVID organ dataset, with stacked bars coloured by cell type. **(F)**, Computational resource usage across datasets, showing peak memory consumption (MB) versus runtime (seconds) on logarithmic scales for model assessment (left) and model selection (right) phases. Rows correspond to sc-COVID multiomics (top), sc-COVID organ (middle), and sc-HTX (bottom). Each point represents an integration method, with point colour and label indicating method identity.

#### 2.4.1 Classification performance on single-cell data integrations

Classification performance across the single-cell case studies showed dataset-specific patterns that differed from those observed in bulk multimodal datasets. Across the three datasets (sc-HTX, sc-COVID multiomics, and sc-COVID multi-organ), the relative ranking of methods varied across datasets (**Fig. 5B1, C1, D1**). In the heart transplant dataset (sc-HTX), RGCCA-N+LDA achieved the highest AUC, followed closely by DIABLO-N and IntegrAO, although overall performance remained moderate across methods (**Fig. 5B1**). In the sc-COVID multiomics dataset, DIABLO-N achieved the highest classification performance, followed by RGCCA-N+LDA and multiview approaches, suggesting stronger discriminative signal in this dataset (**Fig. 5C1**). In the sc-COVID multi-organ dataset, multiview achieved the highest AUC, followed by DIABLO-N and IntegrAO (**Fig. 5D1**). Despite the dataset-specific ranking of methods, DIABLO-N consistently performed well across all three single-cell multiomics datasets, ranking among the top-performing approaches in each case. This suggests that the supervised regularized framework used by DIABLO is robust to the heterogeneous data structures and modality combinations typical of single-cell multimodal studies. In contrast, the performance of other methods varied more substantially between datasets, highlighting that the effectiveness of integration strategies may depend strongly on dataset characteristics such as modality composition, cell-type diversity, and the strength of disease-associated signal captured within each modality.

#### 2.4.2 Biological interpretation for single cell data integration

To evaluate the biological interpretability of features selected by each integration method, we performed pathway enrichment analysis on ranked features derived from each model. The number of significant pathways (FDR<20%) identified by each method varied substantially across datasets (**Fig. 5B2, C2, D2**). In the sc-HTX dataset, several methods identified enriched pathways across multiple cardiac cell types including endothelial cells, cardiomyocytes, macrophages, and fibroblasts (**Fig. 5B2**). RGCCA and DIABLO variants identified the largest number of significant pathways, whereas MOGONET detected the fewest number of enriched pathways. Restricting the pathways to those related to organ transplantation (e.g. adaptive immune system, costimulation by the Cd28 family) showed the RGCCA variants identified the most cell-specific pathways associated with allograft rejection (**Fig. 5B3**). In the sc-COVID multiomics dataset, pathway enrichment was substantially broader, with methods such as MOFA+glmnet, DIABLO-F1, and RGCCA variants identifying large numbers of enriched pathways distributed across immune cell populations including monocytes, neutrophils, B cells, and T cells (**Fig. 5C2**). The associated pathway heatmap demonstrates extensive overlap in SARS-COV2 pathways (e.g. Sars Cov 2 infection, Sars Cov 2 host interactions, potential therapeutics for Sars) across methods (**Fig. 5C3**), consistent with the strong immune activation expected in COVID-19 infection. In the sc-COVID multi-organ dataset, pathway enrichment was concentrated in a subset of immune cell populations, including NK cells, monocytes, and T-cell subsets (**Fig. 5D2**). Notably, a greater number of significant pathways were identified in blood-derived cells compared with airway cells. Among the methods evaluated, MOFA+glmnet, RGCCA variants, and DIABLO-F identified the largest number of enriched pathways. The pathway heatmap indicates that enrichment signals were often modality-specific, with certain pathways consistently identified across methods while others were detected only by individual approaches (**Fig. 5D3**). Together, these results suggest that while many integration methods recover shared biological signals, individual methods may emphasise different aspects of cellular biology depending on how they model cross-modality relationships.

#### 2.4.3 Computational trade-offs in the single-cell setting

Computational scalability differed substantially across integration methods in the single-cell setting (**Fig. 5B4, C4, D4**). Across the three case studies, Multiview was consistently the most time-intensive method, showing the longest runtimes in both model assessment and model selection, especially in the COVID multiomics and COVID multi-organ datasets. IntegrAO was generally the most memory-demanding method, with peak memory often among the highest across panels, and it also showed relatively long runtimes. In contrast, DIABLO was the most computationally efficient overall, with low runtime and low peak memory across all three case studies. MOFA also showed favorable efficiency, remaining relatively fast and memory-efficient, while RGCCA occupied an intermediate position, with modest resource demands overall but somewhat higher runtime or memory in specific datasets. Taken together, these results indicate that DIABLO and MOFA are the most practical methods from a computational perspective, whereas Multiview trades strong performance for substantially higher runtime, and IntegrAO is limited primarily by high memory usage.

## 3 Discussion

MESSI was developed to address a central problem in multimodal integration research: methods are often compared under inconsistent preprocessing, unequal tuning strategies, and non-comparable evaluation schemes, making it difficult to determine whether observed differences reflect true methodological advantages or differences in experimental design. By combining standardized data preparation, interoperable support for R- and Python-based methods, and a nested cross-validation framework for leakage-free model assessment, MESSI provides a reproducible and extensible platform for equitable benchmarking across diverse classes of integration methods.

In the bulk multimodal benchmark, differences in classification performance were generally modest. DIABLO and Multiview tended to rank highest, whereas MOFA+glmnet and MOGONET were weaker overall. These results suggest that predictive accuracy alone may not be sufficient for selecting among integration approaches in many practical settings. Instead, secondary criteria such as interpretability, scalability, and robustness may be equally important. In this regard, our enrichment analyses were particularly informative. DIABLO, RGCCA, MOFA, and IntegrAO consistently identified larger numbers of significant Reactome and oncogenic pathways, and RGCCA, MOFA, and DIABLO most often recovered tissue-relevant PanglaoDB cell-marker signatures. These findings indicate that methods with similar predictive performance can differ meaningfully in the biological plausibility and interpretability of the signals they recover.

The single-cell analyses reinforced the conclusion that method performance is highly dataset dependent. Method rankings varied across the heart transplant, COVID-19 multiomics, and COVID-19 multiorgan datasets, reflecting differences in modality structure, class balance, and feature dimensionality. However, DIABLO performed consistently well across all three datasets, while RGCCA also showed strong performance, particularly in the heart transplant setting. These results suggest that supervised multiblock approaches may be especially robust in high-dimensional pseudobulked single-cell applications. At the same time, pathway enrichment patterns differed across methods, indicating that different integration strategies may emphasize distinct aspects of biology even when their predictive performance is similar.

An important practical finding was that stronger predictive performance did not necessarily require greater computational complexity. DIABLO and RGCCA showed favorable runtime and memory profiles in both bulk and single-cell benchmarks, whereas methods such as multiview cooperative learning and IntegrAO were often more computationally demanding. This is relevant for real-world adoption, particularly in resource-limited environments or in studies involving large-scale single-cell data. Taken together, our results suggest that DIABLO and RGCCA provide a strong balance of prediction, interpretability, and computational efficiency, whereas other methods may be preferable when specific priorities, such as maximizing classification performance or emphasizing particular latent representations, are of primary interest.

This study has several limitations. First, the current benchmark focused on vertical integration and binary classification, and therefore does not yet address other important settings such as survival analysis, regression, clustering, or partially paired mosaic designs. Second, although the benchmark spans a diverse set of bulk and single-cell datasets, it does not capture the full breadth of multimodal biomedical applications. Third, not all available methods could be included, particularly those lacking mature software implementations or those not readily adaptable to a containerized benchmarking workflow. As with any benchmark, there is also an inherent tension between fairness and faithfulness: enforcing a common evaluation framework improves comparability, but may not fully capture the optimal usage of every method.

Overall, MESSI provides a reproducible framework for comparing multimodal integration methods under a common and unbiased model assessment strategy. Our results argue against a universally best method and instead support a task-dependent view of method choice, guided by the balance between predictive performance, biological interpretability, and computational cost. By standardizing benchmarking across heterogeneous tools and data types, MESSI offers a foundation for more transparent method evaluation and for future extensions to broader multimodal learning problems.

## 5 Methods

### 5.1 Integration methods

#### 5.1.1 CaretMultimodal

CaretMultimodal is implemented in R at https://github.com/JoshD898/caretMultimodal. It is built upon caret [50], and caretEnsemble R-package, a late integration approach, where base models are trained independently on each dataset, and their predictions are then aggregated using an ensemble model. We used glmnet with alpha = 0, resulting in ridge regression models.

#### 5.1.3 DIABLO

DIABLO stands for Data Integration Analysis for Biomarker discovery using Latent cOmponents [24]. It extends sparse Generalized Canonical Correlation Analysis (sGCCA) [51] to a supervised setting. Denote 𝑄 normalized, centered and scaled datasets 𝑋^(1)^, 𝑋^(2)^, …, 𝑋^(𝑄)^ such each dataset measures expression levels of 𝑃_1_, …, 𝑃_𝑄_ omics variables on same 𝑁 samples, then sGCCA solves the optimization function for each dimension ℎ = 1, …, 𝐻: maximize

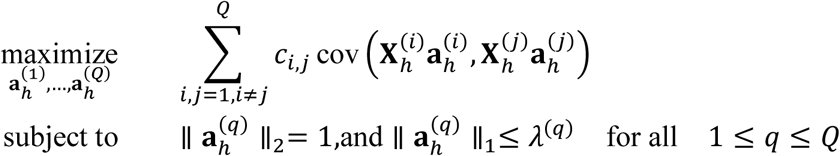

Where:

- 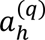 is loading vector on dimension ℎ associated to residual matrix 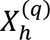 of dataset 𝑋^𝑞^
- 𝐶 = {𝑐_𝑖,𝑗_}_𝑖,𝑗_ is a 𝑄 × 𝑄 design matrix of connection between datasets
- 𝜆^(𝑞)^is a non-negative parameter that controls amount of shrinkage, ultimately number of non-zero coefficients in 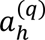.

Then DIABLO extends the above optimization further by substituting one omics dataset 𝑋^(𝑞)^ in above problem with a dummy indicator matrix 𝑌 of 𝑁 × 𝐺 dimension to indicate class membership of each sample, and 𝐺 is number of phenotype or class groups.

#### 5.1.4 IntegrAO

IntegrAO is an unsupervised (and extensible supervised) multi-omics integration framework [52] designed to handle incomplete and partially overlapping multi-omics datasets, produce a unified representation of patients, and enable downstream tasks like patient stratification and classification of new samples even when some omics types are missing.

Denote 𝑀 omics datasets 𝑋^(1)^, 𝑋^(2)^, …, 𝑋^(𝑀)^ with possibly different sets of patients, and let 𝐺_𝑚_ be the patient similarity graph for dataset 𝑋^(𝑚)^. IntegrAO solves the following steps:

Graph construction:

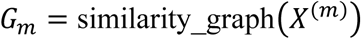

for each modality 𝑚 = 1, …, 𝑀, capturing relationships between patients.

Graph fusion: iteratively update each modality graph using information from the other graphs:

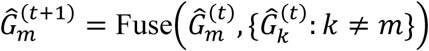

to align shared patient structures across omics.

Patient embeddings via GNN:

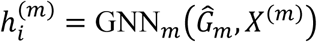

where 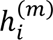 is the embedding of patient 𝑖 from modality 𝑚. Unified embedding:

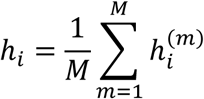

to produce a modality-agnostic representation for each patient.

Optional supervised extension: the unified embeddings can be fine-tuned with class labels to predict phenotypes of new samples, even with missing modalities.

#### 5.1.5 MOFA

MOFA is Multi-Omics Factor Analysis [30]. It can view as a generalization of principal component analysis to multi-omics data. Starting from *M* data matrices *Y* 1*, Y M* of dimensions *N* × *Dm*, where *N* is number of samples and *D_m_* the number of features in data matrix *m*, MOFA decomposes these matrices as:

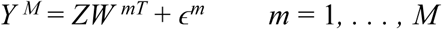

Here, 𝑍 denotes a common factor matrix for all data matrices representing low-dimensional latent variables and 𝑊^𝑚^ as the weight matrices for each data matrix m. And 𝜖^𝑚^ being the omic-specific residual noise term and has different choices of noise model with most frequently used Gaussian noise. MOFA uses variational inference for model fitting and includes automatic relevance determination to promote sparsity in 𝑊^𝑚^. It supports missing values, making it robust for real-world multi-omics datasets with incomplete measurements. Due to the fact that MOFA is unsupervised, hence we fit an additional glmnet model on the 𝑍 factor matrix from MOFA and predict the classes.

#### 5.1.6 MOGONET

MOGONET is named as Multi-Omics Graph cOnvolutional NETworks [31]. It combines graph convolutional network (GCN) for omics specific learning and passed through a view Correlation Discovery Network (VCDN) for multi-omics integration. A different GCN is trained for each omics data type. The loss function for 𝑖 th omics data type GCN_𝑖_ is the following:

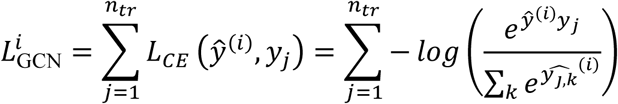

where 𝐿_𝐶𝐸_(.) represents the cross-entropy loss function, 𝑦_𝑗_ is one-hot encoded label of jth training sample, and 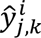 is kth element in vector 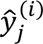.

Furthermore, a VCDN is trained to integrate different omics type by constructing a cross-omics discovery tensor 𝐶_𝑗_. For data with 𝑚 omics data types, each element in 𝐶_𝑗_ can be calculated as:

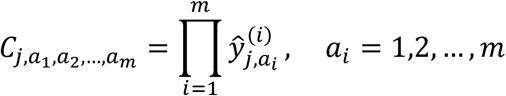

where its loss function is:

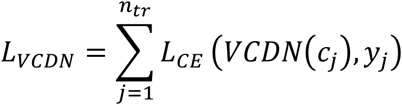

In summary the total loss function of MOGONET could then be summarized as:

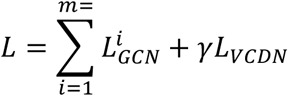

where 𝑚 is number of omics, and 𝛾 is a trade-off parameter between omics-specific classification loss and final classification loss from VCDN.

#### 5.1.2 Multiview

Cooperative Learning (Multiview) is implemented in R with package name multiview [23]. It is a supervised learning framework that integrates multiple data modalities (views) by jointly minimizing prediction error while encouraging agreement across *M* views with the following optimization problem:

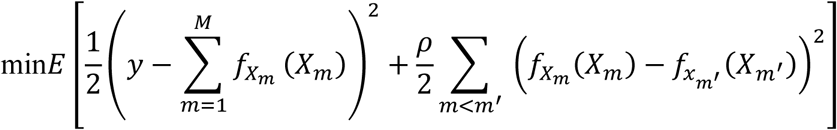

For a hyperparameter *ρ* ≥ 0. The first term of the minimization is the usual prediction error (can also use other loss function, such as mean square error), and the second being an agreement penalty term.

#### 5.1.7 RGCCA

RGCCA is Regularized Generalized Canonical Correlation Analysis [34]. Considering 𝐽 data matrices 𝑋_1_, …, 𝑋_𝐽_, each 𝑛 × 𝑝_𝑗_ data matrix 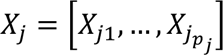 is treated as view. Each view represents set of 𝑝_𝑗_ variables observed on 𝑛 individuals. A core criteria is that individuals has to match across views, where number of variables could differ from one to another. Furthermore, all variables are assumed to be centered.

RGCCA aims to solve the following optimization problem:

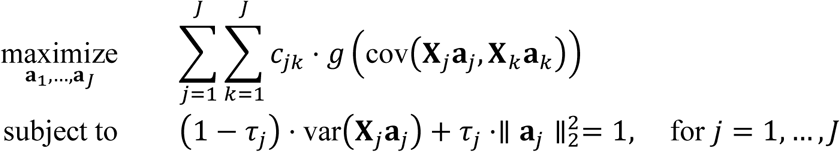

where

- 𝐶_𝑗𝑘_ are elements of the design matrix, indicating the connections between views
- 𝑔 is a continuous convex scheme function applied to the covariance between view components, allowing different optimization criteria like max of sum of covariances or max of sum of absolute values covariances, etc.
- 𝜏_𝑗_ are shrinkage or regularization parameters ranging from 0 to 1.

- 𝜏_𝑗_ = 1 yields maximization of covariance-based criterion, where variance of views dominates over correlation
- 𝜏_𝑗_ = 0 yields maximization of correlation-based criterion, where correlation of connected components is maximized
- 0 < 𝜏_𝑗_ < 1 is a compromise between variance and correlation of the view components

This formulation aims to find weight vectors 𝑎_𝑗_ that maximize the sum of pairwise covariances (or other measures, depending on the choice of 𝑔) between the projected view components 𝑋_𝑗_𝑎_𝑗_

### 5.2 Simulated dataset

In order to benchmark methods under full control and knowledge of the data generating process, we simulated a sets of multiomics data based on the paper [51] with slight modification. We followed A similar approach in the paper mentioned with considering 3 views of omics, i.e. each *n x p_j_* view *X_j_* for *j* = 1, 2, 3 and generated with the following model:

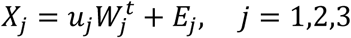

The vector u represents principal components and drawn from multivariate normal distribution:

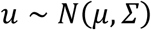

where mu is mean vector with fixed mean for all entries as:

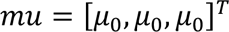

where *µ*_0_ is also controllable as parameter of the package, and default to *µ*_0_ = 0. This is a parameter to help distinguish group differences. The covariance matrix is identity matrix by default representing no correlation across views. We have implemented this simulation in such way to control correlation of each view modifying σ as:

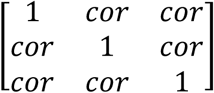

where 𝑐𝑜𝑟 ∈ [0,1], and default to 1. The weights are drawn from the following distribution:

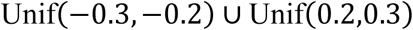

𝐸_𝑗_ is a 𝑛𝑥𝑝_𝑗_ residual matrix drawn from 𝑁(0,1).

### 5.3 Bulk multiomics datasets

We consider multiomics dataset represented by *P* omics matrices *Xi*, where *Xi* is of dimension *n* × *pi* (*n* samples and *p_i_* features) for *i* = 1*, …, P*. The bulk dataset was retrieved from public sources like TCGA from LinkedOmics [53] with R 4.3.3 and libraries TCGAbiolinks 2.30.4 and GEOquery 2.70.0. These datasets were furthered processed to keep matched samples data only (same subjects measured in different omics modalities) Furthermore, they are filtered for omics with nonzero variance during the pipeline execution, as certain methods would not work if zero variance data was present. For a summarized list of real datasets, please refer to **Table 2**. Below we present higher level information of dataset studied in this paper.

#### 5.3.1 GSE38609

This data is retrieved from https://www.ncbi.nlm.nih.gov/geo/query/acc.cgi?acc=GSE38609 with title: *Brain transcriptional and epigenetic associations with the autistic phenotype*. It is composed of an expression data with Illumina HumanHT-12 V4.0 expression beadchip and methylation data with Illumina HumanMethylation27 BeadChip.

#### 5.3.2 TCGA-STES

This data is retrieved from https://www.linkedomics.org/data_download/TCGA-STES/, and studies *Stomach and Esophageal carcinoma* (STES). It is made of methylation data of Illumina HM450K platform, miRNA data of Illumina HiSeq platform, miRgene-level with RPM normalization, reverse phase protein array (protein) gene level data, and RNAseq data normalized counts (Illumina HiSeq platform, Gene-level, RPKM).

#### 5.3.3 TCGA-CHOL

This data is retrieved from https://www.linkedomics.org/data_download/TCGA-CHOL/, and studies *Cholangiocar-cinoma* or commonly known as bile duct cancer. It is composed of an miRNA with RPM normalization, methylation data with Illumina HM450K platform, protein data, and RNAseq data normalized counts (Illumina HiSeq platform, Gene-level, RPKM).

#### 5.3.4 GSE71669

This data is retrieved from https://www.ncbi.nlm.nih.gov/geo/query/acc.cgi?acc=GSE71669 with title: *Integration analysis of three omics data using penalized regression methods: An application to bladder cancer*. It is composed of expression data with Affymetrix Human Gene 1.0 ST Array [transcript (gene) version] and methylation data with Illumina HumanMethylation27 BeadChip (HumanMethylation27_270596_v.1.2).

#### 5.3.5 TCGA-ACC

This data is retrieved from https://www.linkedomics.org/data_download/TCGA-ACC/ and studies *Adrenocortical carcinoma* (ACC). It is composed of methylation data of tumor samples at Gene level from Illumina HM450K platform, miRNA expression for tumor samples with RPM normalization, protein expression normalized (Gene-level), and RNAseq data normalized counts (Illumina HiSeq platform, Gene-level, RPKM).

#### 5.3.6 TCGA-KICH

This data is retrieved from https://www.linkedomics.org/data_download/TCGA-KICH/ and studies *Kidney Chro-mophobe*. It is composed of methylation data of tumor samples at Gene level from Illumina HM450K platform, miRNA expression for tumor samples with RPM normalization, protein expression normalized (Gene-level), and RNAseq data normalized counts (Illumina HiSeq platform, Gene-level, RPKM).

#### 5.3.7 TCGA-MESO

This data is retrieved from https://www.linkedomics.org/data_download/TCGA-MESO/ and studies *Mesothelioma*. It is composed of methylation data of tumor samples at Gene level from Illumina HM450K platform, miRNA expression for tumor samples with RPM normalization, protein expression normalized (Gene-level), and RNAseq data normalized counts (Illumina HiSeq platform, Gene-level, RPKM).

#### 5.3.8 TCGA-SKCM

This data is retrieved from https://www.linkedomics.org/data_download/TCGA-SKCM/ and studies *Skin Cutaneous Melanoma*. It is composed of methylation data of tumor samples at Gene level from Illumina HM450K platform, miRNA expression for tumor samples with RPM normalization, protein expression normalized (Gene-level), and RNAseq data normalized counts (Illumina HiSeq platform, Gene-level, RPKM).

#### 5.3.9 TCGA-BRCA

This data is retrieved from https://www.linkedomics.org/data_download/TCGA-BRCA/ and studies *Breast invasive carcinoma*. It is composed of methylation data of tumor samples at Gene level from Illumina HM450K platform, miRNA expression for tumor samples (Illumina GenomeAnalyzer platform, Normalized, miRgene-level, RPM), protein expression normalized (Gene-level), and RNAseq data normalized counts (Illumina HiSeq platform, Gene-level, RPKM).

#### 5.3.10 TCGA-ESCA

This data is retrieved from https://www.linkedomics.org/data_download/TCGA-ESCA/ and studies *Esophageal carcinoma*. It is composed of methylation data of tumor samples at Gene level from Illumina HM450K platform, miRNA expression for tumor samples with RPM normalization, protein expression normalized (Gene-level), and RNAseq data normalized counts (Illumina HiSeq platform, Gene-level, Normalized log2 RPKM).

#### 5.3.11 TCGA-KIRC

This data is retrieved from https://www.linkedomics.org/data_download/TCGA-KIRC/ and studies *Kidney renal clear cell carcinoma*. It is composed of methylation data of tumor samples at Gene level from Illumina HM450K platform, miRNA expression for tumor samples (Illumina GenomeAnalyzer platform, miRgene-level, Normalized, RPM), protein expression normalized (Gene-level), and RNAseq data normalized counts (Illumina HiSeq platform, Gene-level, RPKM).

#### 5.3.12 TCGA-THCA

This data is retrieved from https://www.linkedomics.org/data_download/TCGA-THCA/ and studies *Thyroid carcinoma*. It is composed of methylation data of tumor samples at Gene level from Illumina HM450K platform, miRNA expression for tumor samples with RPM normalization, protein expression normalized (Gene-level). and RNAseq data normalized counts (Illumina HiSeq platform, Gene-level, Normalized log2 RPKM).

#### 5.3.13 TCGA-BLCA

This data is retrieved from https://www.linkedomics.org/data_download/TCGA-BLCA/ and studies *Bladder urothelial carcinoma*. It is composed of methylation data of tumor samples at Gene level from Illumina HM450K platform, miRNA expression for tumor samples with RPM normalization, protein expression normalized (Gene-level), and RNAseq data normalized counts (Illumina HiSeq platform, Gene-level, Normalized log2 RPKM).

#### 5.3.14 ROSMAP

This data is obtained from AMP-AD Knowledge Portal, and this stands for *The Religious Orders Study and Memory and Aging Project* (ROSMAP) Study, which mainly focus on Alzheimer Disease. It involves genomics, epigenetic, and transcriptomics data.

### 5.4 Multimodal data

#### 5.4.1 Imaging-omics

This dataset was generated by integrating matched spatial transcriptomic, imaging-derived, and cell composition features from kidney tissue regions of interest (http://nanostring-public-share.s3-website-us-west-2.amazonaws.com/GeoScriptHub/KidneyDataset/). Sample annotations were first imported and filtered to retain only regions labeled as “Geometric Segment,” after which unique ROI identifiers were created by combining scan and region labels. The RNA modality was derived from a normalized target count matrix, restricted to geometric segments, relabeled to match the annotation table, log2-transformed, and thresholded so that negative values were set to zero. Imaging features were obtained from precomputed ResNet50 embeddings extracted from ROI images, with filenames parsed to reconstruct identifiers matching the same tissue regions. A third modality, estimated cell-type proportions, was imported from spatial deconvolution output and aligned to the corresponding ROIs using sample identifiers and a cell-type annotation file. All three modalities were perfectly matched across the same regions, with region-level metadata containing disease status as the response variable (diabetic kidney disease, and controls).

#### 5.4.2 Electrical-omics

This dataset was created by combining multimodal measurements from a heart failure cohort that included cellular, electrophysiological, transcriptomic, and proteomic data [46,54]. Demographic and outcome information were imported alongside four feature matrices (https://amritsingh.shinyapps.io/omicsBioAnalytics/): blood cell measurements (cells), Holter monitor–derived variables (holter), mRNA expression (mrna), and protein abundance (proteins). Because the files were already aligned by subject, a common sample identifier was assigned across all modalities by creating matched row names for each dataset. The clinical outcome used for prediction was hospitalization status.

#### 5.4.3 Clinical-omics

This dataset was constructed by integrating matched clinical and omics measurements from the P1000 prostate cancer cohort[47]. Copy-number alteration (CNA) data, RNA expression data (TPM), clinical annotations, and a binary response file were first imported from the processed study files (https://zenodo.org/records/10775529). Samples were then harmonized across all four sources by retaining only subjects present in every dataset. To reduce missingness in the clinical modality, a subset of informative clinical variables was selected, including age, mutation-related measures, sequencing coverage, fraction of genome altered, ploidy, purity, and sample type, and only samples with complete data for these variables were kept. The final outcome label included two categories (primary or metastasis), while the remaining clinical variables were used as predictors. The resulting matched dataset therefore contains three aligned modalities per subject, CNA, RNA, and clinical features.

### 5.5 Single-cell multiomics data

#### 5.5.1 COVID19-multiomics single-cell transcriptomics

This dataset was generated from a published single-cell multiomics study[55] of the peripheral immune response in severe COVID-19 and built by integrating donor-level pseudobulk RNA and protein measurements derived from the same single-cell dataset. A Seurat object downloaded from the COVID-19 Cell Atlas (https://www.covid19cellatlas.org/index.patient.html) was first imported, and separate `SingleCellExperiment` objects were created for the RNA assay and the imputed protein assay. For each modality, cells were annotated with donor identity and coarse cell type, then aggregated using pseudobulking so that expression values were summarized within each donor-by-cell-type combination. For RNA, raw counts were summed across cells, combined across cell types, normalized using TMM, transformed with `voom`, and filtered to retain genes with mean raw count greater than one. For proteins, imputed protein values were averaged within each donor-by-cell-type combination and similarly reshaped into a donor-level feature matrix. In both modalities, the donor-by-cell-type matrices were reorganized so that each donor became one sample and each feature represented a specific cell-type–feature pair. Donor-level metadata were then extracted from the original single-cell object, including COVID-19 status as the response variable. The final harmonized dataset was stored as a `MultiAssayExperiment` containing matched RNA and protein assays for each donor, together with links to the original publication and source atlas.

#### 5.5.2 COVID19-multiorgan single-cell transcriptomics

This dataset was constructed by integrating pseudobulk single-cell transcriptomic profiles from two tissue compartments, airway and peripheral blood mononuclear cells (PBMCs), from a COVID-19 cohort[56]. Separate `h5ad` objects were first imported for airway and PBMC samples (https://www.covid19cellatlas.org/), and each dataset was restricted to relevant case-control groups and annotated with donor or patient identifiers and detailed cell-type labels. Within each compartment, cells were aggregated by donor-by-cell-type using pseudobulking, summing expression counts across cells for each cell type within each individual. The resulting donor-by-cell-type count matrices were then normalized with `TMM` and transformed using `voom`, after which low-abundance genes were filtered out. For both airway and PBMC data, these normalized matrices were reshaped so that each donor became a single sample and each feature corresponded to a specific cell-type–gene pair. Donors shared between the airway and PBMC datasets were then intersected to create matched multimodal profiles across both compartments. Finally, donor-level clinical metadata were aligned to the shared donors and used to define the outcome variable (COVID19, and healthy), and the airway and PBMC pseudobulk matrices were stored together in a `MultiAssayExperiment`, yielding a harmonized single-cell multimodal dataset for COVID-19 analysis.

#### 5.5.3 Heart transplant single-cell spatial transcriptomics

This dataset (GSE290577) was created from spatial transcriptomic Xenium data from heart transplant biopsies[57] by converting single-cell counts into a cell-type-resolved pseudobulk multimodal object. First, the Xenium object and accompanying sample metadata were loaded, and the analysis was restricted to pre-biopsy samples while excluding mixed rejection cases. Cell-level metadata were then aligned to the selected samples and used to define the sample identifier, patient grouping variable, rejection response label, and cell-type annotation. Raw RNA counts were extracted from the Xenium object and used to build a `SingleCellExperiment`, after which counts were aggregated with muscat by cell type within each sample to generate pseudobulk expression matrices. Each cell-type-specific pseudobulk assay was then normalized separately using TMM followed by log-CPM transformation, while samples with zero library size were retained as all-zero columns to preserve cohort structure. For each cell type, genes with zero variance across samples were removed, and the remaining normalized matrix was stored as a separate `SummarizedExperiment`. These cell-type-specific assays were then assembled into a `MultiAssayExperiment`, with a sample map linking all experiments to the same biopsy-level metadata. Finally, experiments were filtered to retain only cell types with at least 15 nonzero samples in every rejection category, producing a harmonized cell-type-resolved `MultiAssayExperiment` suitable for comparing no rejection, with rejection across heart transplant biopsies.

## Data and code availability

All data generated or analyzed in this study are publicly available in Zenodo under accession 10.5281/zenodo.18842128. The MESSI Nextflow pipeline is available on GitHub at CompBio-Lab/MESSI-pipeline, and the scripts used for the MESSI manuscript are available at CompBio-Lab/MESSI_manuscript.

